# Accommodation and wavelength: the effect of longitudinal chromatic aberration on the stimulus-response curve

**DOI:** 10.1101/2023.06.20.545755

**Authors:** Maydel Fernandez-Alonso, Abigail P. Finch, Gordon D. Love, Jenny C. A. Read

**Affiliations:** Biosciences Institute, Newcastle University, UK; Now at Translational Sensory & Circadian Neuroscience Group, Max Planck Institute for Biological Cybernetics, Germany; Department of Physics, Durham University, UK; Department of Computer Sciences, Durham University, UK; Now at School of Computing, University of Leeds, UK

**Keywords:** accommodation, longitudinal chromatic aberration, stimulus-response curve, pupil size

## Abstract

The longitudinal chromatic aberration (LCA) of the eye creates a chromatic blur on the retina that is an important cue for accommodation. While this mechanism can work optimally in broadband illuminants such as daylight, it is not clear how the system responds to the narrowband illuminants used by many modern displays. Here, we measured pupil and accommodative responses as well as visual acuity under narrowband LED illuminants of different peak wavelengths. Observers were able to accommodate under narrowband light and compensate for the LCA of the eye, with no difference in the variability of the steady-state accommodation response between narrowband and broadband illuminants. Intriguingly, our subjects compensated more fully for LCA at nearer distances. That is, the difference in accommodation to different wavelengths became larger when the object was placed nearer the observer, causing the slope of the accommodation response curve to become shallower for shorter wavelengths and steeper for longer ones. Within the accommodative range of observers, accommodative errors were small and visual acuity normal. When comparing between illuminants, when accommodation was accurate, visual acuity was worst for blue narrowband light. This cannot be due to the sparser spacing for S-cones, since our stimuli had equal luminance and thus activated M-cones roughly equally. It is likely because ocular LCA changes more rapidly at shorter wavelength, and so the finite spectral bandwidth of LEDs corresponds to a greater dioptric range at shorter wavelengths. This effect disappears for larger accommodative errors, due to the increased depth-of-focus of the eye.

## Introduction

The purpose of the ocular lens is to adjust the optical power of the eye so as to produce a sharp, in-focus image on the retina. However, its ability to achieve this is affected by the eye’s longitudinal chromatic aberration (henceforth LCA). The refractive index of the eye decreases with an increase in wavelength, such that for a broadband light, the shorter wavelengths come into focus in front of the retina and the longer wavelengths behind the retina. The resulting defocus as a function of wavelength is shown in Figure 1. The total defocus across the entire visible spectrum is approximately 2 dioptres. This means that the lens is unable to simultaneously optimise ocular power for all visible wavelengths. If green light is in focus, as shown in Figure 1, red and blue light will be out of focus, with positive and negative defocus error respectively.

**Figure 1.**
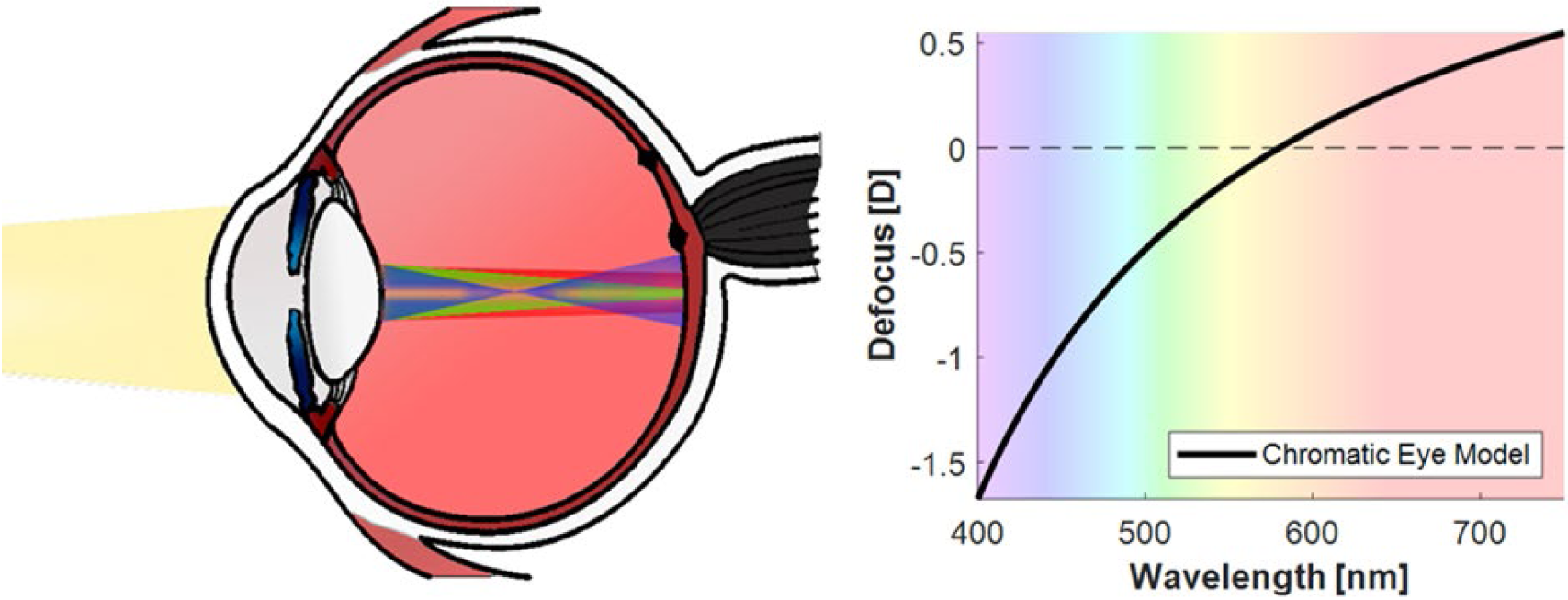
The longitudinal chromatic aberration of the eye. Diagram showing the change in refractive index with wavelength (left), and the defocus caused by LCA as a function of wavelength according to the chromatic eye model (right). The chromatic eye model specifies the eye’s refractive error as D(λ) = p − q/(λ − c), where λ is the wavelength of light in micrometres and D(λ) is the refractive error in dioptres. For the three parameters, we took the values used by Marimont & Wandell (1994): p = 1.7312, q = 0.63346, c = 0.21410, and the reference wavelength that is kept in-focus is 580 nm

### Role of LCA

Thus, LCA has several implications for visual perception in broadband polychromatic light, such as daylight. Most obviously, image quality can be reduced in polychromatic light compared to monochromatic light (Campbell & Gubisch, 1967; Aggarwala et al., 1995). Similarly, contrast sensitivity is greater if chromatic aberration is corrected with achromatic lenses or reduced by using monochromatic light (Yoon & Williams, 2002; Williams et al., 2000; Artal et al., 2010).

However, LCA also implies a greater depth of field in polychromatic light, that is, a greater range of accommodation values for which the image will appear acceptably sharp in the retina, since at least one wavelength is in focus (Campbell, 1957; Campbell & Gubisch, 1967; Marcos et al., 1999). An increased depth of field in polychromatic light has been proposed as a possible explanation for the non-linearity of the human accommodation function. A steady-state error is typically found when accommodation is measured for different distances with white light, and it has been proposed that this could be explained by a change in the component wavelength that is brought into focus at different distances (Ivanoff, 1949). For nearer distances, short wavelength components would be brought into focus, while for further distances, long wavelength components would be the ones in focus. Due to the increased depth of field, the image formed in the retina would remain acceptably sharp in both conditions. However, evidence already exists against this idea (Bobier et al., 1992; Labhishetty et al., 2021).

In addition to reducing retinal image contrast and increasing the depth of field, LCA can also contribute an odd-error cue to accommodation. In an eye free of aberrations, positive and negative defocus both produce the same effect on the point-spread function at a given wavelength. Thus in monochromatic light, it is impossible to know whether image contrast will be improved by reducing or increasing ocular power. However, in polychromatic light, one can infer the sign of defocus by comparing the amount of blur at different wavelengths. If red light is blurred more than blue, accommodation should be relaxed, and vice versa. There is evidence that this polychromatic blur serves as an important cue to accommodation (Fincham, 1951; Kruger et al., 1993).

### Narrow-band primaries

One of the main differences between digital displays and the natural environment is in the spectral distribution of the light they emit or reflect. While daylight is composed of a smooth spectrum and natural objects tend to have broad spectral reflectance functions (Krinov, 1953),Click or tap here to enter text. most digital displays take advantage of the fact that human vision is trichromatic and make use of only three lights or primaries to show us different images. These primaries – red, green, and blue– give rise to a spectral distribution with multiple narrowband peaks rather than a smooth spectrum, with modern displays increasingly making use of particularly narrowband light sources such as Light Emitting Diodes (LEDs), organic LEDs (OLEDs) and lasers. As these lights differ significantly from the natural light the human visual system evolved to accommodate under, it is important to understand how they affect the accommodative response of the eye in order to maximise the quality of the image perceived in these displays.

Narrowband primaries might in particular, affect the way the visual system makes use of the longitudinal chromatic aberration (LCA) of the eye to aid accommodation. This blur would be significantly reduced when accommodating under the individual narrowband primaries of a display. Furthermore, LCA would cause a shift in the best-focus distance for each individual wavelength, so observers would need to adjust their response accordingly.

#### Do narrow-band primaries affect the accuracy of accommodation?

Both reduced spectral bandwidth and removing LCA can negatively impact the accuracy of the dynamic accommodation response of the eye. Kruger et al. (1993) measured accommodation responses to a sinusoidally moving target illuminated by either white broadband light or a light of 10 nm spectral bandwidth, while the LCA of the eye was either normal, removed, or reversed. They found that accommodative gain decreased and phase-lag increased, when LCA was neutralised or when the target was illuminated by narrowband light. Furthermore, reversing the sign of LCA severely disrupted the accommodation response of observers and their ability to track the object while moving in depth. In a later study, Aggarwala et al. (1995) used a sinusoidally moving target illuminated by lights of 10 nm, 40 nm and 80 nm of spectral bandwidth, as well as a broadband white light. Their results indicated that as the spectral bandwidth of the light increased, accommodative gain increased and phase lag decreased, with the broadband white light enabling significantly more accurate dynamic responses than the 80 nm spectral bandwidth light. These authors performed another similar study where the sinusoidally moving target was illuminated either by one of ten narrowband lights of 10 nm spectral bandwidth and peak wavelengths between 430nm and 670nm that were viewed through an achromatizing lens, or by a white broadband light that was viewed with and without the achromatizing lens (Aggarwala, Nowbotsing, et al., 1995). They found that accommodative gain tended to be higher and phase lag lower when the target was illuminated by white light with LCA intact; however, their results also indicated that there was great inter-subject variability in the accommodative responses to the stimuli between observers, as some of the subjects seemed to be able to track some of the monochromatic targets moving in depth and accommodate to them reasonably well. The authors concluded that narrowband illumination was a poor stimulus for accommodation and suggested that visual displays that used narrowband primaries were likely to reduce the ability of the eye to maintain accurate focus.

Other studies have not found evidence that the absence of LCA has a detrimental effect on accommodative responses, particularly when the targets are stationary. Bobier et al. (1992) measured the accommodation stimulus-response curve for a broadband target when the LCA of the eye was normal, neutralized, increased, and reversed. They found that the slopes of the accommodation functions did not change in any of the six subjects for any of the conditions tested, with only one participant showing an effect on the reversed LCA condition, with a lower intercept and steeper slope. Thus, it seems that neither removing or increasing LCA had a significant effect on the static accommodation response of participants, and even when the sign of LCA was reversed, participants were able to maintain their steady-state accommodation responses. When looking at the variability of the responses under broadband and narrowband illumination for stationary targets, Atchison et al (2004) found that none of their five observers had more intra-trial variability in their accommodation when looking at targets illuminated by narrowband red or blue light, than when observing targets under broadband light, suggesting that they were able to maintain focus just as well under reduced spectral bandwidth.

The differences between these sets of studies could suggest that LCA is more important for dynamic accommodation than for steady-state responses. That is, the visual system uses LCA to detect when a change in accommodation is required, as well as the direction of the change, but once it is focused on a target, it can maintain accommodation via other cues or mechanisms. Kotulak et al. (1995) found evidence that this might be the case. They measured dynamic and steady-state responses to stimuli of varying spectral bandwidth and found that increasing bandwidth caused an increase in the gain of dynamic responses (although no differences in phase lag), but that it had no effect on the steady-state error of accommodative responses. However, later studies by Kruger at al. (1997) provided evidence contrary to this. Participants viewed stationary square-wave gratings placed at distances of 0, 2.5 and 5 dioptres, and under three conditions of illumination: light of 10 nm spectral bandwidth, broadband light with normal LCA, and broadband light with reverse LCA. They found that all subjects accommodated accurately in the normal LCA condition, 38% had difficulty maintaining focus in monochromatic light at near and far (5 and 0 dioptres), and 88% could not maintain focus at both near and far when LCA was reversed. They speculated that the detrimental effects of reduced spectral bandwidth on the steady-state accommodation response were only detectable at distances that were far away from the tonic state of accommodation, and that this could be the reason for the differing findings in previous studies, as those had used distances that were nearer the resting accommodative state of the eye. Thus, LCA could also be an important cue for steady-state accommodation responses, and the reduced spectral bandwidth of narrowband primaries in a display could impair the accuracy of this response, particularly at near and far distances.

#### Do observers adjust for LCA when accommodating to different primaries?

When accommodating to monochromatic or narrowband stimuli of different wavelengths, the accommodative response shifts in the direction predicted by LCA, that is, higher accommodation for longer wavelengths and lower for shorter ones (Charman, 1989; Donohoo, 1985; Lovasik & Kergoat, 1988b, 1988a; Seidemann & Schaeffel, 2002), although the magnitude of the dioptric shift has not always been up to the magnitude predicted by the LCA defocus (Donohoo, 1985).

However, these studies have usually tested targets placed at only one or two fixed distances. Few studies to date have looked at the effect that narrowband light of different wavelengths has on the overall accommodation stimulus-response curve. Charman & Tucker (1978) measured the accommodation of seven subjects at multiple target vergences for white light and for different narrowband illuminants. Most of their participants were experienced observers and were able to accommodate under monochromatic light as accurately as under broadband light; however, their one naïve observer was not initially able to accommodate to the narrowband targets, requiring further training to be able to maintain accommodation for these stimuli. They also found that there was a dioptric shift in the accommodation responses of participants with wavelength, but no difference in their accuracy, such that the stimulus-response curves of one subject showed similar lags and leads for all colours tested. They did find however, that for blue light some observers had a slightly shallower slope, which they attributed to a combination of a small increase in LCA with accommodation (approximately 3% per dioptre of accommodation), as well as reduced acuity for blue light in some subjects.

More recently, Jaskulski et al. (2016) measured the subjective depth of field of seven subjects for targets at distances of 0, 2 and 4 dioptres, and when illuminated under broadband and monochromatic red, green and blue light. The measurements were performed in the paralyzed eye, while the higher order aberrations of the accommodated eye of each participant were simulated using an adaptive optics system. They found that the slopes of the best focus position as a function of accommodative demand were lower than one, but similar between monochromatic and white light. Furthermore, they found no significant differences in the subjective depth of field under different monochromatic lights, and the depth of field for white light was greater at all distances by approximately 14%, although the differences were not statistically significant.

There are some limitations in these two studies that should be considered. Firstly, they both used mostly well-trained observers with experience in accommodation experiments, as described by the authors. We have seen so far that there can be significant inter-subject variability in the responses to monochromatic stimuli or to broadband stimuli when LCA has been removed or reversed (Aggarwala, Nowbotsing, et al., 1995), which can explain some of the contradictory findings in the literature; and naïve observers can struggle to accommodate in monochromatic light without receiving training beforehand (Charman & Tucker, 1978). This means that these findings might not be representative of the general population or the average untrained user of visual displays. Furthermore, Jaskulski et al. (2016) paralyzed the accommodative and pupil response of the eye and estimated accommodation from the subjective reports of perceived blur from the observers, which might not be a good indication of their real accommodative responses with natural pupil sizes.

### The present study

The literature reviewed so far shows that there are not yet clear answers to the two questions posed above. As well as being of interest scientifically, these questions are important given the increasing use of narrow-band primaries in modern visual displays. Any effect on the accuracy or precision of accommodation could lower image quality and increase the risk of visual fatigue. Thus, we aimed to address both questions together, using a larger sample with a greater proportion of untrained observers. Furthermore, we allowed the accommodation and pupil size of observers to vary freely, to increase the ecological validity of the results and more closely match a real-life scenario of subjects viewing images in a digital display with narrowband primaries. Finally, we also concurrently measured visual acuity using a staircase procedure to explore the impact that any difference in accommodation to narrowband stimuli might have on the ability of subjects to resolve small targets, when compared to accommodation in broadband light where LCA is available as a cue.

## Methods

In experiments 1 and 2, the accommodation function was sampled by changing the physical distance of the stimuli, with the angular size of the diffuser changing concurrently in experiment 1 and being kept constant in experiment 2. In experiment 3, the accommodation function was measured by using trial lenses to simulate a larger range of distances, and the visual acuity of participants was measured concurrently. Experiments 1 and 2 used the same apparatus, thus, they are described together, while experiment 3 is described separately where necessary.

### Participants

Participants were recruited from students, staff, and the external pool of participants of the Biosciences Institute of Newcastle University for experiments 1 and 2, and only from students and staff of the Institute of Biosciences for experiment 3. The study adhered to the Declaration of Helsinki, was approved by Newcastle University Faculty of Medical Sciences Research Ethics Committee and written consent was obtained from each subject. Table 1 shows the breakdown of the included sample for each individual experiment.

**Table 1.** Sample description for experiments 1, 2 and 3.

|  | **Participants** | **Mean age (± SD)** | **Sex** |
| --- | --- | --- | --- |
| Experiment 1 | 8 | 26.5 (± 2.5) | 4 females, 4 males |
| Experiment 2 | 9 | 25.9 (± 2.5) | 4 females, 5 males |
| Experiment 3 | 10 | 29.5 (± 2.4) | 6 females, 4 males |

### Apparatus: Experiments 1 & 2

The stimulus consisted of a Maltese cross printed on a transparent film and placed on top of a diffuser, which was mounted on a box containing six LEDs and centred to the right eye. The box was placed on a 2.5m long rail positioned at the height of participants’ eyes, which allowed to change the physical distance of the stimulus. The stimulus was presented at six distances between 3D and 0.5D in steps of 0.5D (corresponding to metric distances of 33.3cm, 40cm, 50cm, 66.7cm, 100cm, and 200cm).

For experiment 1, we kept constant the physical size of the diffuser (8.5 by 8.5cm) and the Maltese cross (5 by 5cm) across the different distances, thus changing its angular size. The angular size of the diffuser changed between 14.5° and 2.4° in steps of 2.4° for the different distances, while the angular size of the Maltese cross changed between 8.6° and 1.4° in steps of 1.4°. For experiment 2, we kept the angular size of the diffuser and the Maltese cross constant across the different distances at 2.5° and 1.5° respectively.

The refractive state of the eye and the pupil diameter was measured dynamically at 50 Hz using a photorefractor with pupillometry capabilities (the PowerRef 3 from PlusOptix, Nuremberg, Germany; Plusoptix.com). Two Arduino Uno boards controlled the stimuli and were connected to the photorefractor to synchronise the recordings with the stimuli. A representation of the experimental setup is shown in Figure 2.

**Figure 2.**
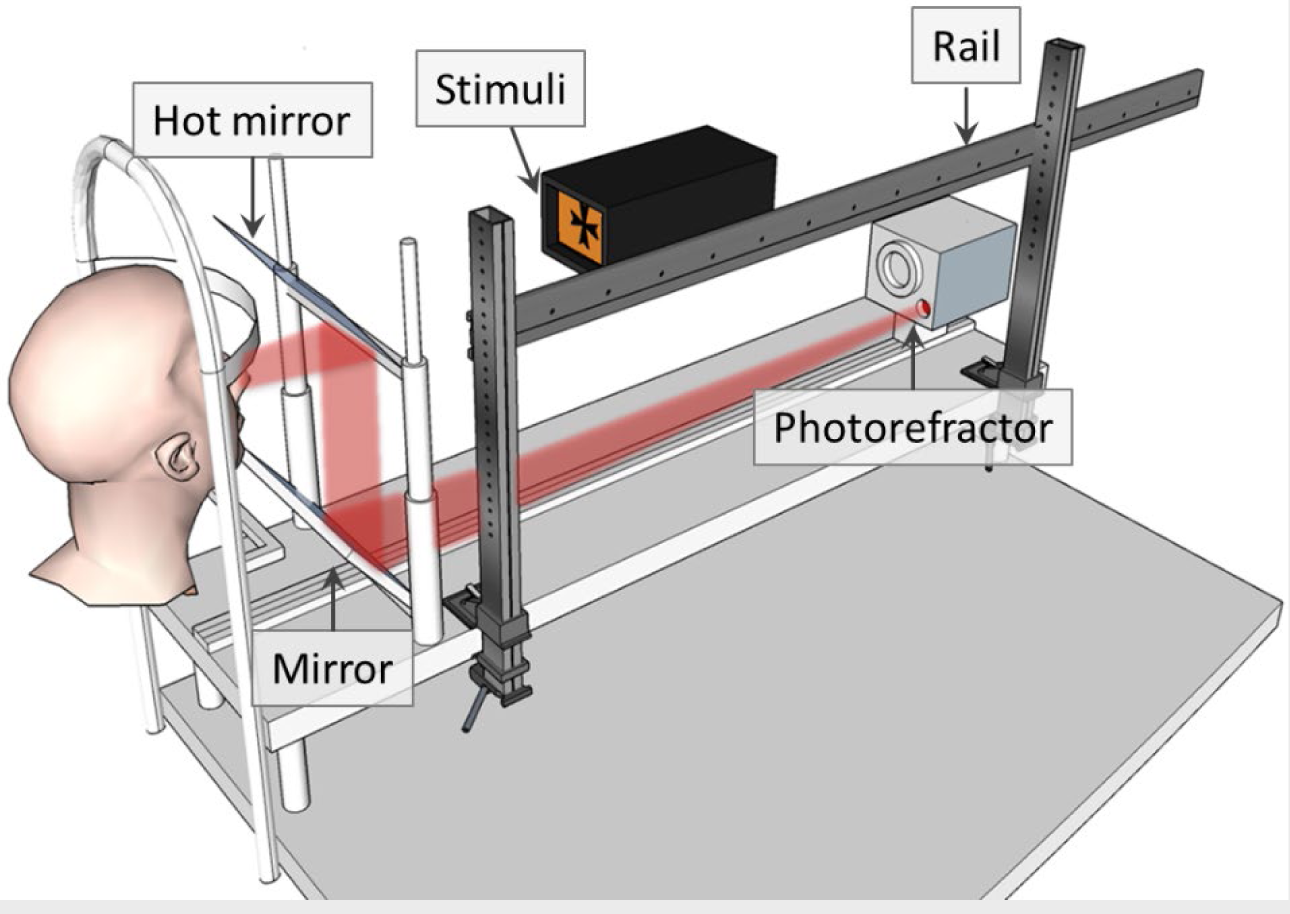
Diagram of the experimental setup for Experiments 1 and 2. The stimuli consist of a black Maltese cross on a bright background formed by a diffuser back-illuminated by LEDs. The colour of the background varies depending on which LEDs are used. The physical distance of the stimuli is varied by moving them along a rail. The observer’s accommodative state is monitored using a photorefractor, which views the observer’s eyes via a hot mirror, that transmits visible light while reflecting infra-red.

The different spectra were created using six LEDs, five of which were narrowband and one white LED, with the latter being combined with the narrowband LEDs to create a broadband spectral distribution that approximated a D65 illuminant (see Figure 3). A driver circuit was built for each of the LEDs, and their luminance was controlled through pulse-width modulation from the Arduino Uno boards (at a frequency of 980Hz). The circuit was designed such that the luminance of the LEDs varied minimally over time, by increasing the resistance and decreasing the current through each LED. During the first 10 seconds after each LED was turned on, the luminance remained constant for all LEDs except the red one, for which luminance decreased by ∼0.7 cd/m^2^. Radiance measurements of the LEDs were taken with a CS-2000 Konica Minolta spectroradiometer at different duty cycles and over time. We multiplied each radiance spectral distribution by the CIE physiologically-relevant luminous efficiency function V(λ) (Stockman et al., 2008) to obtain the peak wavelength and full width at half maximum (FWHM, see Figure 2). We integrated these to obtain the luminance, which we confirmed was a linear function of duty cycle for each LED. During the experiment, the luminance of all stimuli was kept constant at 10cd/m^2^.

**Figure 3.**
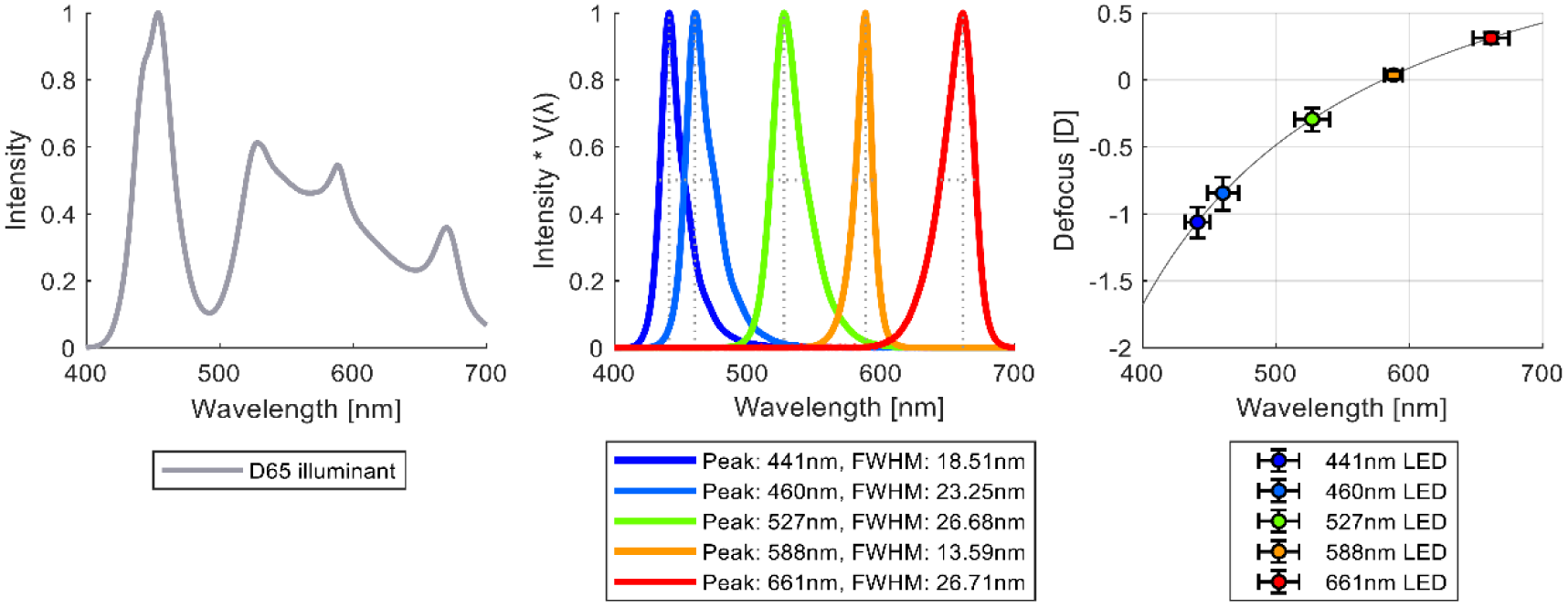
Normalised spectral distributions of the D65 broadband illuminant (left) and the narrowband LEDs (middle), and the defocus caused by LCA for the peak wavelengths of the LEDs (right), with horizontal error bars representing the FWHM and vertical error bars the corresponding spread in defocus.

### Apparatus: Experiment 3

The stimulus consisted of different Landolt C figures that were presented in an active-matrix organic light emitting diode (AMOLED) screen placed at a fixed distance of 1m (1D). The screen had a size of 6.84cm by 12.2cm, and a resolution of 1080 by 1920 pixels, and was from a OnePlus 3T mobile phone device.

To simulate the defocus caused by viewing the stimuli at different distances, 9 trial lenses were used with powers that ranged from -2D to 7D in steps of 1D. The stimuli were viewed through the lenses, which were placed over the right eye in light-tight goggles. The left eye was covered by a 720nm infrared filter that occluded the visual stimuli while allowing the refractive state and pupil diameter of the eye to be measured by the PowerRef 3 photorefractor. A graphical representation and photos of the experimental setup are shown in Figure 4.

**Figure 4.**
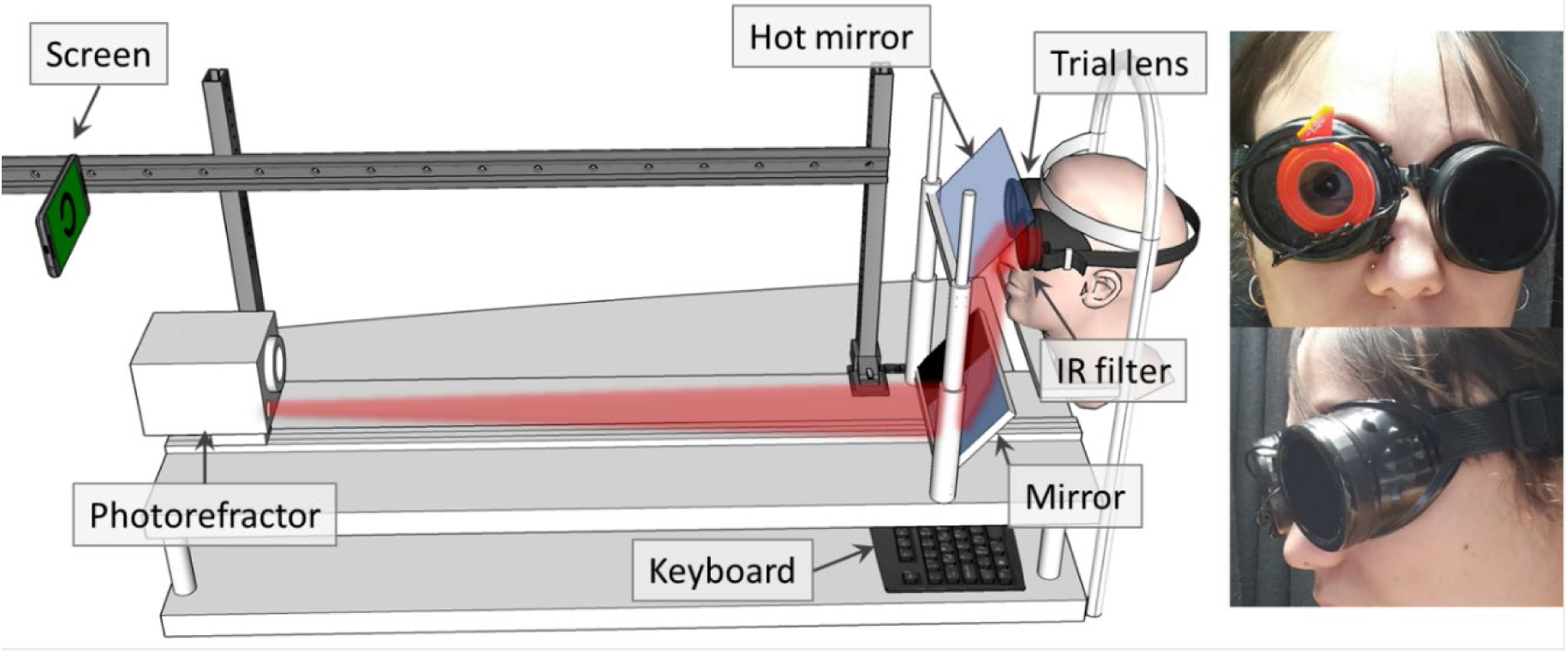
Representation and photos of the experimental setup for Experiment 3. The set-up is similar to that shown in Figure 1, except now the stimuli are presented on an AMOLED screen at a fixed distance of 1m. The observer views the stimuli monocularly through a lens placed over their right eye. The left eye is covered by a filter which blocks visible light, while allowing the photorefractometer to monitor refractive state and pupil size using infra-red.

The AMOLED screen and experimental routine were controlled from a computer running MATLAB (The MathWorks Inc., 2019), which was also connected to the photorefractor to synchronize the stimuli being presented with the recordings. The Landolt C figures were dynamically created using the Psychophysics Toolbox (Kleiner et al., 2007). Figure 5 shows the spectral distributions of the screen primaries, as well as the defocus caused by the LCA of the eye for their peak wavelengths (Thibos et al., 1992).

**Figure 5.**
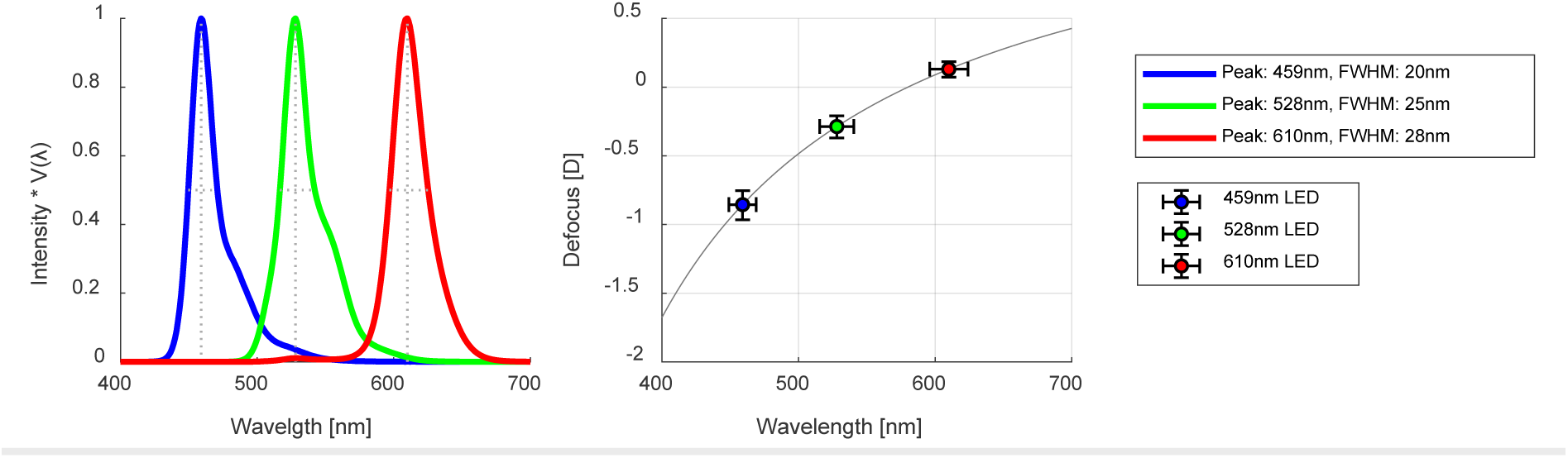
Normalised spectral distributions of the screen LED primaries (left), and the defocus caused by LCA for their peak wavelength (right), with horizontal error bars representing the FWHM and vertical error bars the corresponding spread in defocus.

Radiance measurements of the screen primaries were taken with a CS-2000 Konica Minolta spectroradiometer at different intensities and over time. The peak wavelength and luminance of the LEDs were calculated using the CIE physiologically-relevant luminous efficiency function (Stockman et al., 2008). During the experiment, the three primaries of the screen were used at a fixed luminance of 15cd/m^2^ when used on their own to give narrowband illumination, and when they were combined to create a broadband illumination, each primary was given a luminance of 5cd/m^2^ for the same total luminance of 15cd/m^2^.

### Photorefractor calibration

The photorefractor used in these experiments (PowerRef 3 from Plusoptix) consists of an array of nine infrared LEDs (850nm) located eccentrically below an infrared camera that records at 50Hz. Two mirrors reflect the infrared light into the retina (see Figure 3), which diffusely reflects this light back into the camera, and depending on the refractive state of the eye, the light reflected will vary in intensity vertically across the pupil. An inbuilt calibration factor then converts this slope of intensity across the pupil into a defocus value that indicates the refractive state of the eye.

This slope-based eccentric infrared (IR) photorefraction offers a convenient non-invasive way to measure refraction dynamically over a large range of dioptric values (-7D to 5D from the camera position at 1D) and pupil sizes (∼3mm-8mm); however, the accuracy of the results will largely depend on the calibration factor, which is often obtained from a sample of mostly Caucasian individuals. Previous studies have demonstrated that ethnic differences (Sravani et al., 2015) and further inter-individual differences (Bharadwaj et al., 2013; Ghahghaei et al., 2019) affect this calibration factor, reducing the accuracy of the results. They have also suggested how a correction factor specific to each individual can be quickly found and used to reduce these errors significantly, that is independent of the viewing distance used To find this individual correction factor we followed the method described by Sravani et al. (2015). Further details about its implementation are given in the supplements.

### Design and procedure: Experiments 1 & 2

At the start of the experimental session, participants read the information sheet and signed the consent form. Their visual acuity was then measured at near and far distances using a Snellen chart and a logMAR (logarithm of the Minimum Angle of Resolution) test, respectively. All participants had a visual acuity of logMAR 0.25 or better without the need for spectacle or contact lenses. That is, they could read characters that were smaller than 8.9 arcmin wide with a stroke width of 1.8 arcmin. The photorefractor calibration procedure was then performed.

During the experiment, their left eye was covered using an eyepatch and they sat with their head placed on a chinrest. They were instructed to fixate on the stimulus presented and to keep it in focus with as much effort as if they were reading a book. A button placed next to them allowed them to pause the task at any time, and frequent breaks were given throughout the experiment.

The distance of the stimuli was varied between experimental blocks, with the order of the distances being randomised between participants. In experiment 1, the size of the diffuser and fixation was kept constant, while in experiment 2, it was changed according to the distance of the target to keep a constant angular size. Within each experimental block, the target was illuminated by the five narrowband illuminants and the broadband illuminant, with their order being randomised. In experiment 1, each illuminant was presented for 8 seconds and repeated at least five times at each of the six distances, for a total of 180 trials. In experiment 2, each illuminant was presented for 3 seconds and repeated 12 times each at each of the six distances, for a total of 432 trials. Between trials, the target was illuminated in both experiments with the orange (588nm) LED to keep a constant luminance adaptation and to start at a relatively similar accommodation value before the target stimuli was presented. Both experiments took approximately one hour to complete.

### Design and procedure: Experiment 3

At the start of the experimental session, participants read the information sheet and signed the consent form. The photorefractor calibration procedure was then performed, and they were then given instructions for the visual acuity task. During the experiment, participants sat with their head placed on a chinrest, while wearing a pair of light-tight goggles that had an infrared filter over the left eye and allowed us to place different trial lenses over the right eye. Frequent breaks were given between experimental blocks and participants could pause the experiment at any time.

To measure visual acuity, we used a 4 Alternative Forced Choice (4-AFC) task with a best PEST staircase procedure of 24 trials (Kingdom & Prins, 2016). Each Landolt C was presented until the participant gave an answer, and the entire staircase procedure took between 20 and 30 seconds to complete. The background of the Landolt C targets was varied for each staircase according to the four illuminants used in the experiment (three narrowband and one broadband). The order of the illuminant was randomised within each experimental block, and a break of 5 seconds was given between each where no stimuli was presented. For each experimental block, a different trial lens was placed in front of the participant’s right eye to add different values of defocus to the stimulus, and the order of the lenses was randomised between participants.

The distance of the stimuli in dioptres was calculated as a function of the physical distance of the screen in dioptres (*P_scrn_*), the power of the different lenses placed in front of the eye (*P_scrn_*), and the distance from the eye to the lens (*x_lens_*), such that:

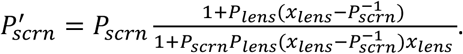

Furthermore, the visual acuity thresholds obtained in degrees of visual angle were corrected for the small magnification the lenses produced, which was calculated as:

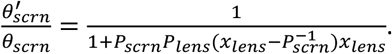

The corrected thresholds in degrees of visual angle were then transformed to logMAR units by converting the values into minutes of visual angle and calculating the base-10 logarithm.

### Data processing and analysis

To analyse the refractive and pupil size recordings, the data points where the pupil was not found were identified as blinks and excluded, as well as 60ms before and 120ms after each blink. Blinks would on occasion cause big spikes in the refractive data, thus, any data points where reported refraction was greater than 25D were also excluded. To allow time for the participants to accommodate, the first 1500ms of refractive and pupil size data in each trial were excluded from further analysis in experiments 1 and 2. Similarly, the first 2000ms of data in each trial were excluded in experiment 3. Finally, any trial with less than 1000ms of measurements in experiments 1 and 2, or 2000ms of measurements in experiment 3 were excluded as well. The calibration correction factor obtained for each participant was then applied to the refraction data, and the median accommodation and pupil size was obtained for each trial.

To perform the analysis on the slopes of the accommodation function, we first determined the linear portion of the accommodation response curve. For this, we calculated the gradient of the accommodation response for each illuminant at each distance, as well as the median gradient, and at any distances where the gradient decreased by 50% or more when compared to the median, the response was deemed to be saturated. These results were visually inspected, and some manual corrections were performed, although they mostly agreed well with the visual evaluation of the experimenter.

For the slope and within-trial response variability analyses (sections 2.3.2 and 2.3.4), the data of experiment 3 was divided into smaller subsets to improve the fit results. The trials in this experiment had a duration of between 20 to 30 seconds, so each one was divided in equal subsets of at least 5 seconds of duration. For all other analyses (i.e., sections 2.3.3, 2.3.5 and 2.3.6) the data was not divided.

Several linear mixed models were used to analyse the effect of distance and illuminant on the slope of the accommodation function (section 2.3.2), the effects of distance on the difference in accommodation to different wavelengths (section 2.3.3), the effects of illuminant and accommodation on response variability (section 2.3.4), the effects of the effects of accommodation on pupil diameter (section 2.3.5), and the effects of accommodative error on visual acuity (section 2.3.6). In all cases, we used the maximal random-effects structure without convergence issues. All models were fitted with the maximum likelihood estimation method, and all fits and corresponding residuals were visually inspected to verify that all assumptions were met. For the slope analyses (section 2.3.2), the median refraction data within each experiment was weighted by the number of valid measurements obtained within each trial as a proportion of the total number of measurements possible. That is, trials where fewer refractive measurements were obtained due to blinking or other factors, were assigned a lower weight in the model fits.

The data processing and most of the model fits were performed using MATLAB, while the model fits on visual acuity performed in section 2.3.6 and the post-hoc analyses were done using R (R Core Team, 2021), particularly the lme4 library (Bates et al., 2023) and the emmeans library (Lenth et al., 2023).

## Results

Figure 6 shows a typical accommodative response for the different illuminants used in experiments 1 and 2. As shown, a change in the refractive state of the eye occurs after approximately 300ms from stimuli presentation, alongside pupil constriction for some of the illuminants presented. After 1000ms, the refractive state of the eye remains relatively constant, while the pupil size slowly increases. For subsequent analysis, we use the steady-state response defined as the median value after the initial exclusion period (see Methods).

**Figure 6.**
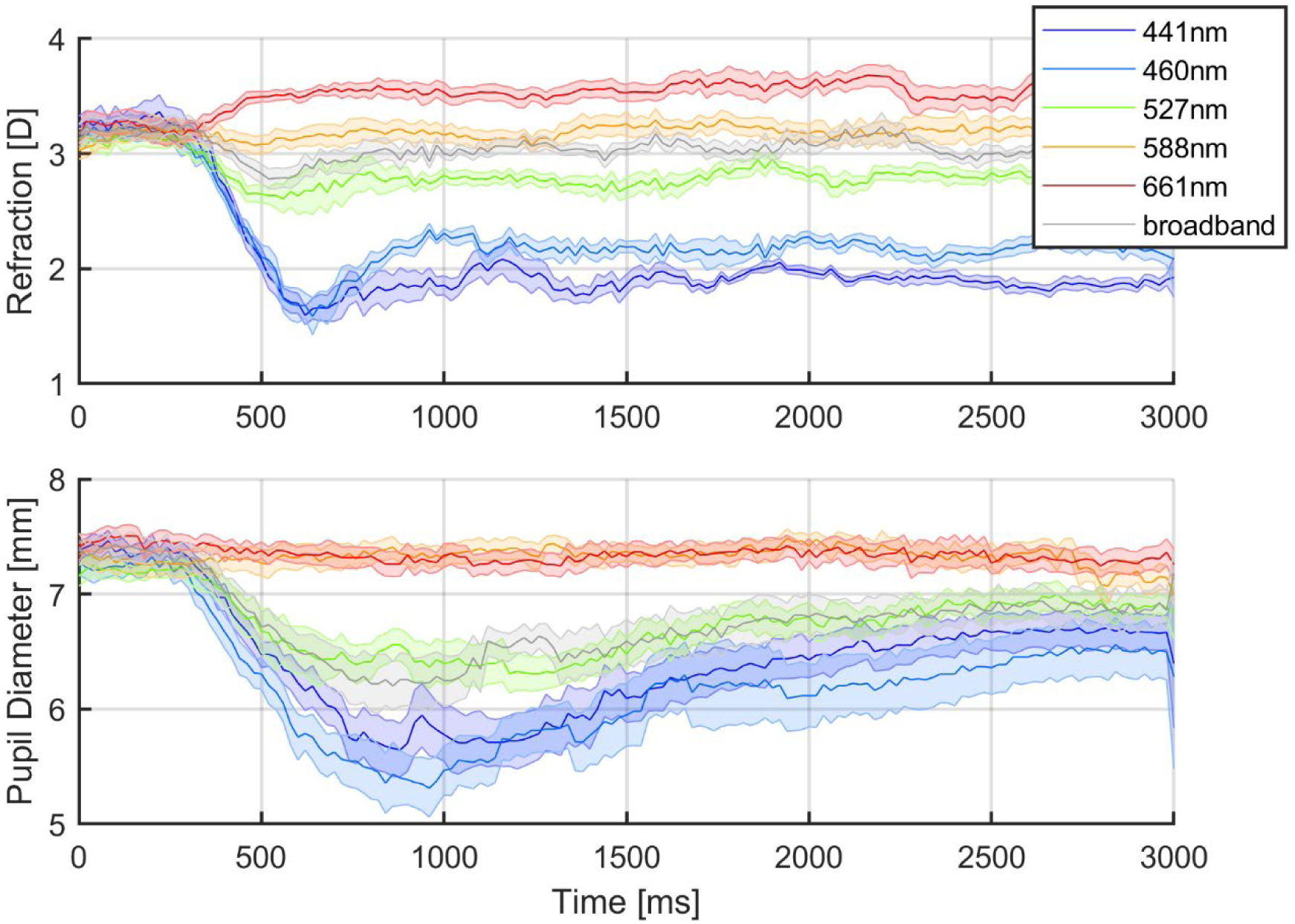
Average accommodation response of one participant to different illuminants presented at 33.3cm (3 dioptres) and repeated 12 times each. The top panel shows the mean refractive state of the eye and the bottom panel the mean pupil diameter as a function of time. The continuous lines represent the mean, and the shaded areas represent the standard error of the mean. The time point of 0ms represents the start of the trials, when the illuminant changed from the 588nm (orange) to the corresponding illuminant as indicated by the legend. This figure was generated by Matlab script fit_z_accTraces_e1.m in the code repository.

### Effects of LCA on the accommodation response curve

An example plot of steady-state response is shown in 7. This shows the mean +/- SEM of the median response on each trial, for the different illuminants. For this subject, the minimum refraction is around 1.3D; stimuli further than this are out of accommodative range. Within the accommodative range of the subject, the response for a given illuminant is a quasi-linear function of distance, with the absolute value of accommodation changing in accordance with the defocus caused by LCA for each illuminant (i.e., at the same distance, observers accommodate less for shorter wavelengths and more for longer ones). An interesting feature of the data, found in most subjects, is that the slope of the response seems to change for each illuminant, with a shallower slope observed for shorter wavelengths and a steeper slope for longer ones.

**Figure 7.**
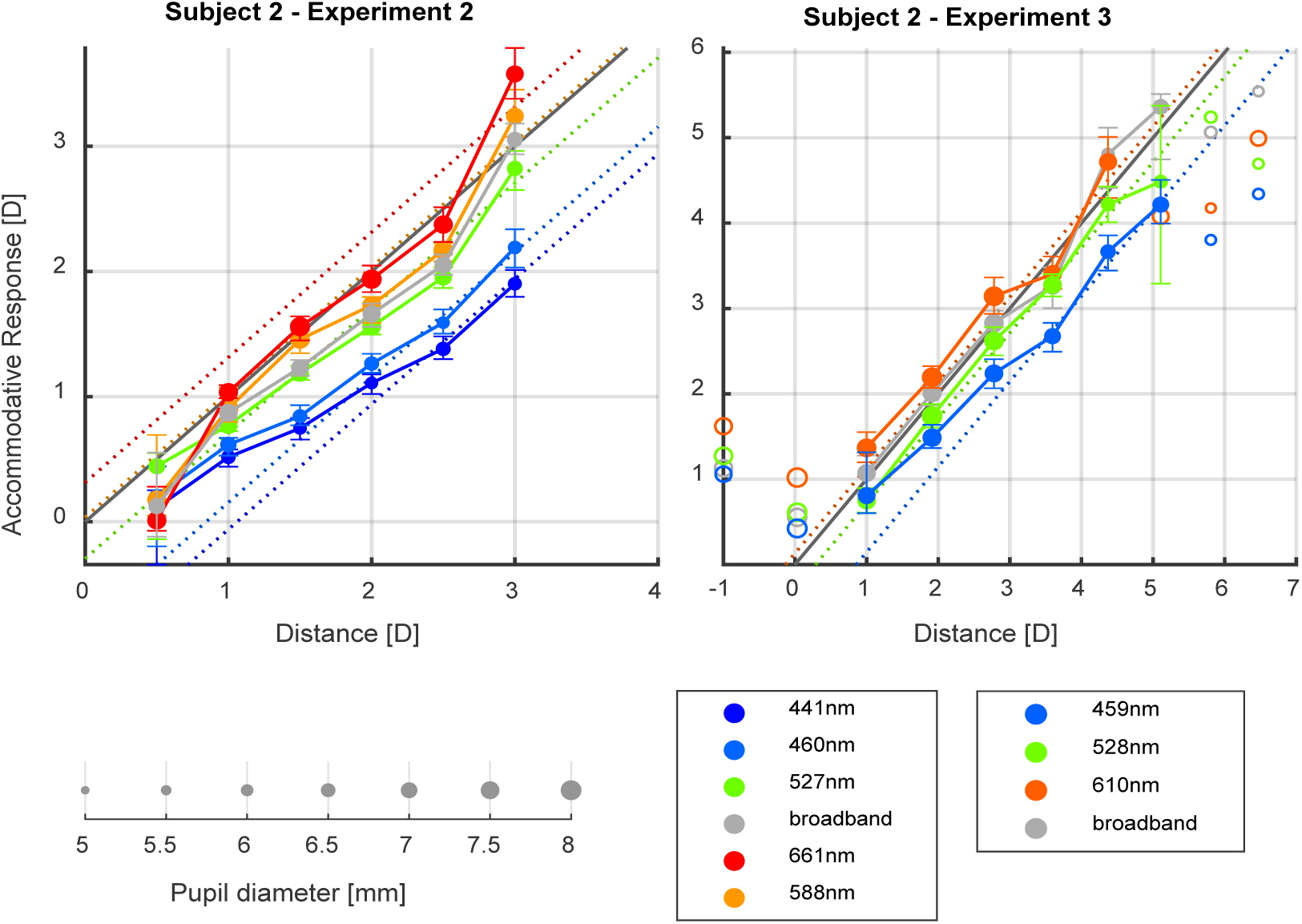
Stimulus-response function for one example participant who participated in Experiments 1 and 3. Median ocular refraction is plotted as a function of the stimulus distance, both in dioptres, for the different illuminants. Filled symbols linked with lines show points classified as being on the linear portion of the stimulus-response curve; empty symbols show points classified as outside the range of accommodation and thus not used for fitting. Symbol size represents median pupil diameter.

To quantify this effect, we first determined the linear portion of the stimulus-response curve and excluded the stimuli that fell outside the accommodative range. Further details about this procedure are given in the supplements. The accommodation response curves of individual participants for each illuminant in the three experiments are also provided in online data repository.

We fitted linear mixed-effects models, since these allow us to obtain slope and intercept estimates for each illuminant, as well as account for individual differences between observers. The fits were performed on the median accommodation response and only over the linear portion of the accommodation response curve. Three participants of experiment 3 (subjects 15, 21 and 22) were excluded from this analysis, as their response curves for most illuminants were only linear over 2 or 3 distances.

For each experiment, the linear mixed models were fitted with predictors of distance in dioptres, illuminant, and their interaction, and random intercepts and slopes of participant (i.e., the effect of distance, illuminant and their interaction was allowed to vary randomly among observers). Illuminant was used as a categorical predictor because the broadband illuminant with no peak wavelength was included, and because the change in slope with peak wavelength for the narrowband illuminants might not be linear (since the defocus caused by LCA as a function of wavelength is not linear). These models were compared in each case with a simpler model that contained no interaction term between distance and illuminant, so the effect of wavelength on accommodation would be constant regardless of distance and the slope for all illuminants would be the same. Results from the Likelihood Ratio Test (LRT) and the Akaike information criterion (AIC) comparison indicated that the model with the interaction term fitted the data better and had greater predictive power in all cases: for experiment 1 (χ^2^ (55) = 177.6, *p*<0.001, ΔAIC= 67.6), experiment 2 (χ^2^ (55) = 265.8, *p*<0.001, ΔAIC= 155.8), and experiment 3 (χ^2^ (24) = 146.92, *p*<0.001, ΔAIC= 98.9).

The results of the linear mixed models are illustrated in Figure 8 and given in full in the supplemental materials. The individual slopes and intercepts estimated for each subject (obtained from the estimated coefficients of the random effects of the model) are presented in the supplemental materials (Supplementary Table 6).

**Figure 8.**
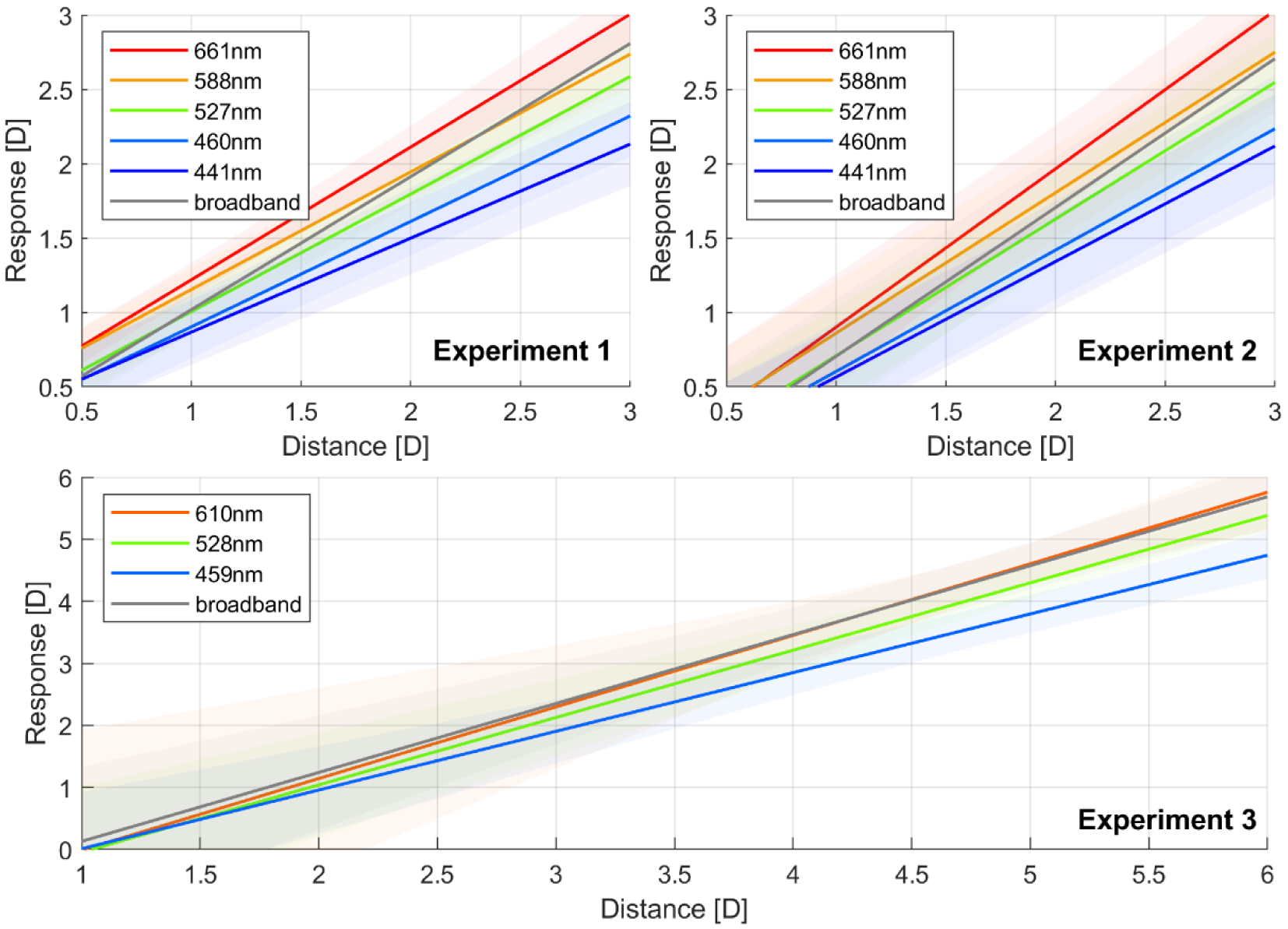
Estimated refraction as a function of distance for the linear portion of the accommodation response curve of all participants in each experiment. The continuous line represents the estimated response and the shaded areas the 95% confidence intervals. The different colours represent the different illuminants used. This figure can be generated with fig_b_oneLMM_123_wDistance_accResp.m in the code repository, and it uses the shadedE rrorBar function by Campbell (2023).

The results show, as illustrated in Figure 8, that the slope of the accommodation response curve for narrowband illuminants becomes shallower as the peak wavelength decreases. Thus, the accommodation responses to narrowband illuminants are mostly similar at optical infinity, but as the stimulus nears the observer, the difference in accommodation to different wavelengths increases in correspondence with the defocus caused by LCA. This results in steeper slopes for longer wavelength illuminants, and shallower slopes for shorter peak wavelengths.

To illustrate how the change in slope for different illuminants affect the lags and leads of the linear portion of the accommodation response curve, we show in Figure 9 the accommodative error as a function of distance for each illuminant. We calculated the accommodative error by subtracting the demand from the predicted response, thus, a negative error indicates the eye is under-accommodating or focusing farther away than where the target is (accommodative lag), and a positive error indicates that the eye is focusing nearer than the stimulus (accommodative lead). The accommodative demand is given by the distance of the stimuli and the defocus caused by LCA for the peak wavelength of the narrowband illuminant, which we calculated following the chromatic eye model by (Thibos et al., 1992). As illustrated, the increased difference in the accommodative response to different wavelengths as the stimulus is placed nearer, corresponds with the change in demand caused by LCA. In other words, participants are increasingly compensating for LCA as the target is placed at nearer distances, causing the accommodative error to become both smaller and less dependent on wavelength. Furthermore, accommodation is more accurate for middle wavelengths over most distances, with a tendency to overaccommodate for shorter wavelengths and under-accommodate for longer ones, and the accommodative errors for all wavelengths and in all three experiments seem to approach a small negative value of approximately -0.5D rather than zero, indicating a small accommodative lag at nearer distances. This lag may in fact maximise image quality, due to factors such as spherical aberration (Labhishetty et al., 2021).

**Figure 9.**
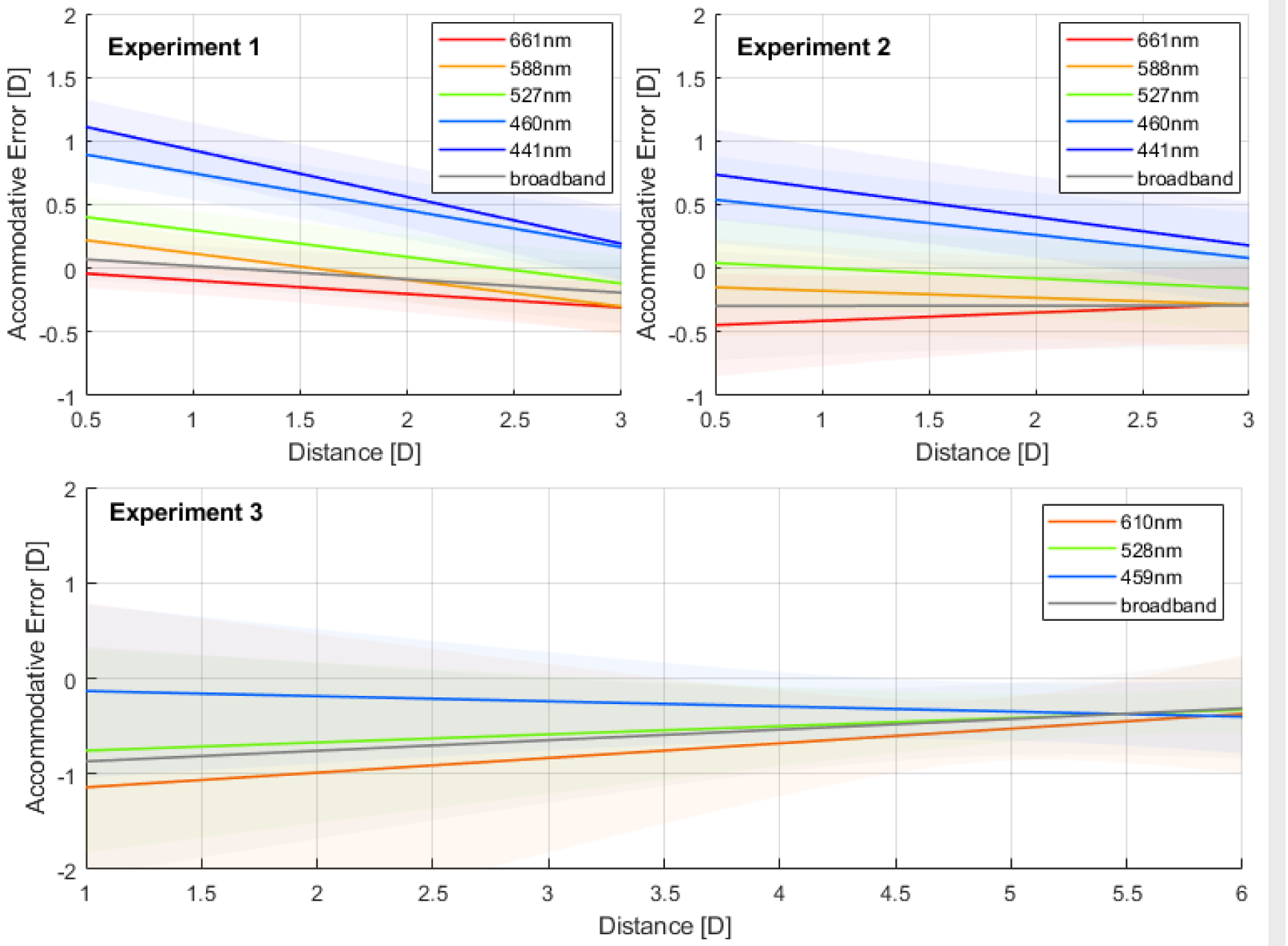
Accommodative error as a function of distance for different wavelengths, as predicted by the linear mixed effects models fitted to the data of each experiment. The continuous lines of different colours represent the predicted responses for different illuminants, as indicated by the legend, and the shaded regions represent the 95% confidence intervals. Note the wider range of the horizontal axis for Experiment 3. This figure can be generated with fig_b_LMM_123_wDistance_accError.m in the code repository and it uses the shadedErrorBar function by Campbell (2023).

### Effects of distance on the accommodation response to different wavelengths

As shown previously, the extent to which participants change their accommodative responses under illuminants of different wavelengths to compensate for the LCA of the eye, changes with the distance of the stimulus. To determine the rate of this change, we fitted a linear mixed model on the relative difference in accommodation between wavelengths, with predictors of distance in dioptres, the defocus predicted by the Chromatic Eye model, and their interaction, and random slopes and intercept of participant. The accommodation responses were centred around the response to the green illuminant (527nm in experiments 1 and 2, 528nm in experiment 3) for each subject at each distance and the defocus predicted by the Chromatic Eye model was set to be zero at 527.5nm.

While in the previous analysis, illuminant was being treated as a categorical predictor, and no assumptions were being made about how it affected accommodation, here we are using the LCA defocus predicted by the chromatic eye model for the peak wavelength of the narrowband illuminants and examining how well it predicts the differences in accommodation between illuminants. A slope of 1 for this relationship would indicate that observers are fully compensating for the LCA of the eye when accommodating to the narrowband illuminants, while a slope of 0 would indicate that there are no differences in the accommodation response to different wavelengths (i.e., they are not correcting for the defocus caused by LCA). Furthermore, we are also exploring here how this relationship between LCA defocus and the difference in accommodation to different wavelengths changes as a function of distance. Based on the previous results presented thus far, we would expect nearer distances to cause an increase in the effect that LCA has on the accommodation response.

The results of the fitted linear mixed model are shown in Table 2. A model that included experiment as a fixed effect as well as its interaction with distance and defocus was also fitted, however no significant effect of experiment was found, and a LRT and AIC model comparisons revealed that the model including experiment as a factor was not significantly better than the simpler model (ΔAIC = - 45.85; χ^2^ (8) = 5.26, *p* = 0.730). This means that the differences in the design of the three experiments did not influence the extent to which participants correct for LCA, once the individual differences between participants had been accounted for. Other fits including the median pupil diameter of participants as an interacting factor were also attempted, however this variable was not found to have a significant effect.

**Table 2.** Linear mixed model results of accommodation to different wavelengths relative to the green illuminant, as a function of distance, the defocus caused by LCA, and their interaction. Coefficient estimates, their 95% confidence interval (CI 95%), and the random effects standard deviations (RE SD) are shown, as well as the t-tests results with degrees of freedom (df), t-ratios, and p-values. Parameters in bold italics are significant at the 0.05 level.

| Parameter | Estimate | CI 95% | t-ratio | df | p-value | RE SD |
| --- | --- | --- | --- | --- | --- | --- |
| Intercept | 0.05 | -0.03 0.14 | 1.19 | 5205 | 0.233 | 0.19 |
| Distance [D] | -0.02 | -0.05 0.00 | -1.71 | 5205 | 0.088 | 0.05 |
| LCA Defocus [D] | -0.26 | -0.76 0.23 | -1.04 | 5205 | 0.296 | 1.14 |
| Distance [D] * LCA Defocus [D] | 0.28 | 0.08 0.47 | 2.82 | 5205 | 0.005 | 0.45 |

As seen in Table 2, the results agree with the previous slope estimations, with the effect of LCA on accommodation increasing by a factor of 0.28 for every dioptre of increase in the distance of the stimulus (95% CI between 0.08 and 0.47, *t*(5205) = 2.82, p = 0.005). This means that at a distance of approximately 4.5 dioptres, participants change their accommodation responses to compensate for the defocus caused by LCA to the full extent predicted by the Chromatic Eye model. Furthermore, we see that at a distance of 0 dioptres, participants do not significantly change their accommodation to different wavelengths (*t*(5202)= 1.04, *p*=0.296), albeit there is considerable variability between subjects at this distance, as indicated by the large standard deviation of the random effects and the wide confidence intervals. Finally, as the predictor was centred at 527.5nm and the response was centred by the 527nm and 528nm illuminants, effectively removing the effect of distance, we see that distance has no significant effect on accommodation when LCA defocus is 0 (i.e., at 527.5nm).

An illustration of the results of the model plotted as a function of wavelength and at different distances is shown in Figure 10 (left), as well as the fitted responses of two subjects (middle and right). As observed, the confidence intervals are wider at the distance of 0.5 dioptres, reflecting the uncertainty of the predictions likely caused by the inter-observer differences being greater at this distance. This is illustrated in the differences between subject 3 and subject 8, as the latter shifted their accommodative responses to correct for LCA to a greater extent than the former when the stimulus was placed at 0.5 dioptres. However, we see that for nearer distances, their responses are more similar. In summary, distance had a significant effect on the dioptric shift observed in the accommodative responses of participants to narrowband illuminants of different wavelengths, however there was considerable inter-subject variability, particularly at further distances.

**Figure 10.**
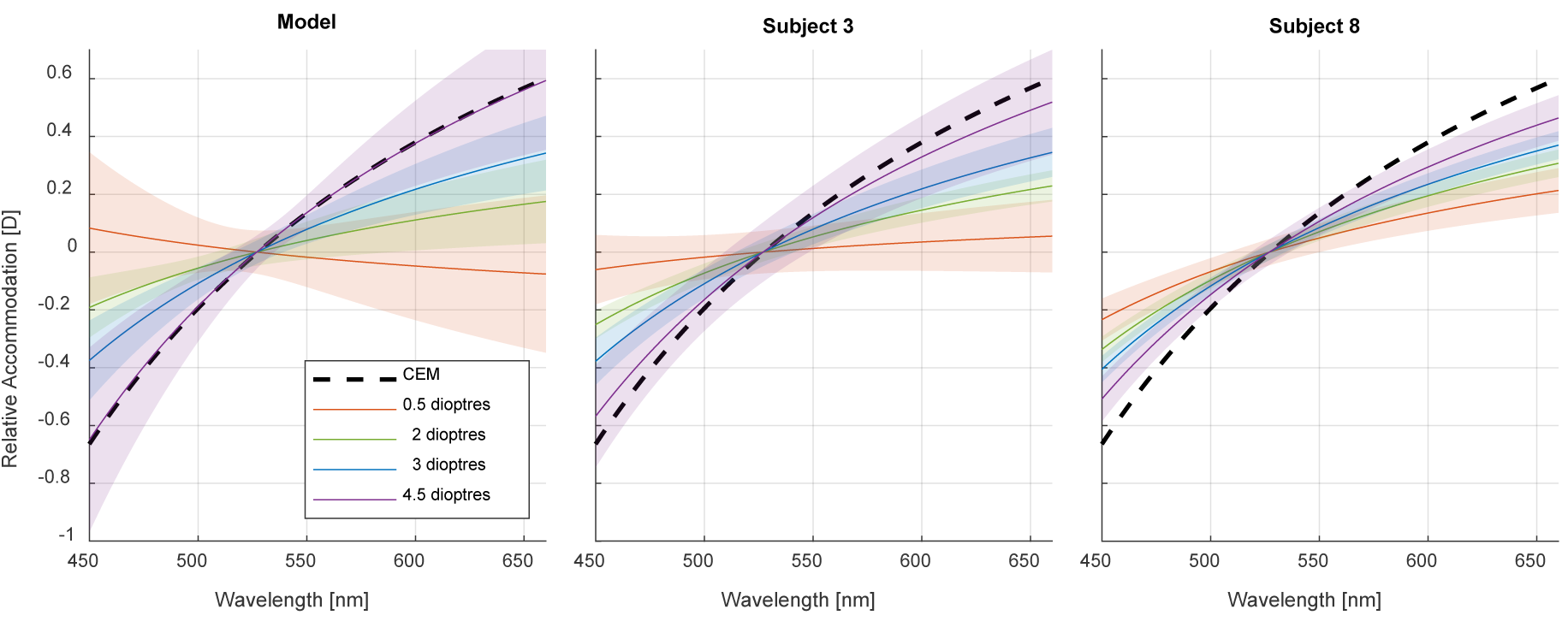
Relative changes in accommodation to different wavelengths as predicted by the linear mixed model (left), and for subjects 3 and 8 (middle and right). The response at each distance was centred by the 527 and 528nm illuminants, so it represents the relative difference in accommodation to these wavelengths. The black dashed line represents the defocus caused by LCA as predicted by the Chromatic Eye model (and centred at 527.5nm). The continuous coloured lines represent different distances as indicated by the legend, and the shaded regions represent the corresponding 95% confidence intervals. This figure can be generated with the fig_i_LCAwithDistance.m file in the code repository, and it uses the shadedErrorBar function by Campbell (2023)

### Variability in the accommodation responses to narrowband and broadband illuminants

In our experiments we recorded the refraction of the eye dynamically at a frequency of 50Hz, which allows us to assess the within-trial variability of the steady-state accommodation response over time for the different illuminants used. In other words, once participants accommodate to a target, how much does the response fluctuate over time, and are there any differences between narrowband and broadband illuminants, and between narrowband illuminants of different wavelengths? Furthermore, as fluctuations in accommodation are known to increase with increasing accommodative power, we also evaluated the effect of the mean refractive state as a predictor.

To obtain a measurement of intra-trial accommodation variability, we fitted a linear function through the refractive response measured in each trial as a function of time and obtained the root-mean-squared errors (RMSE). This approach has been used previously for similar purposes (MacKenzie et al., 2010) and has the advantage of penalizing larger fluctuations in accommodation more and maintaining the units of the response. In Figure 11 we illustrate the distributions of the RMSEs as a function of mean accommodation and illuminant. Since the same illuminants were used in experiments 1 and 2, the data obtained in both was combined. As shown, the within-trial variability seems to increase with increasing mean accommodative state, as well as appearing to be higher for shorter-wavelength illuminants than longer-wavelength ones, with no obvious differences observed between the latter and the broadband illuminants.

**Figure 11.**
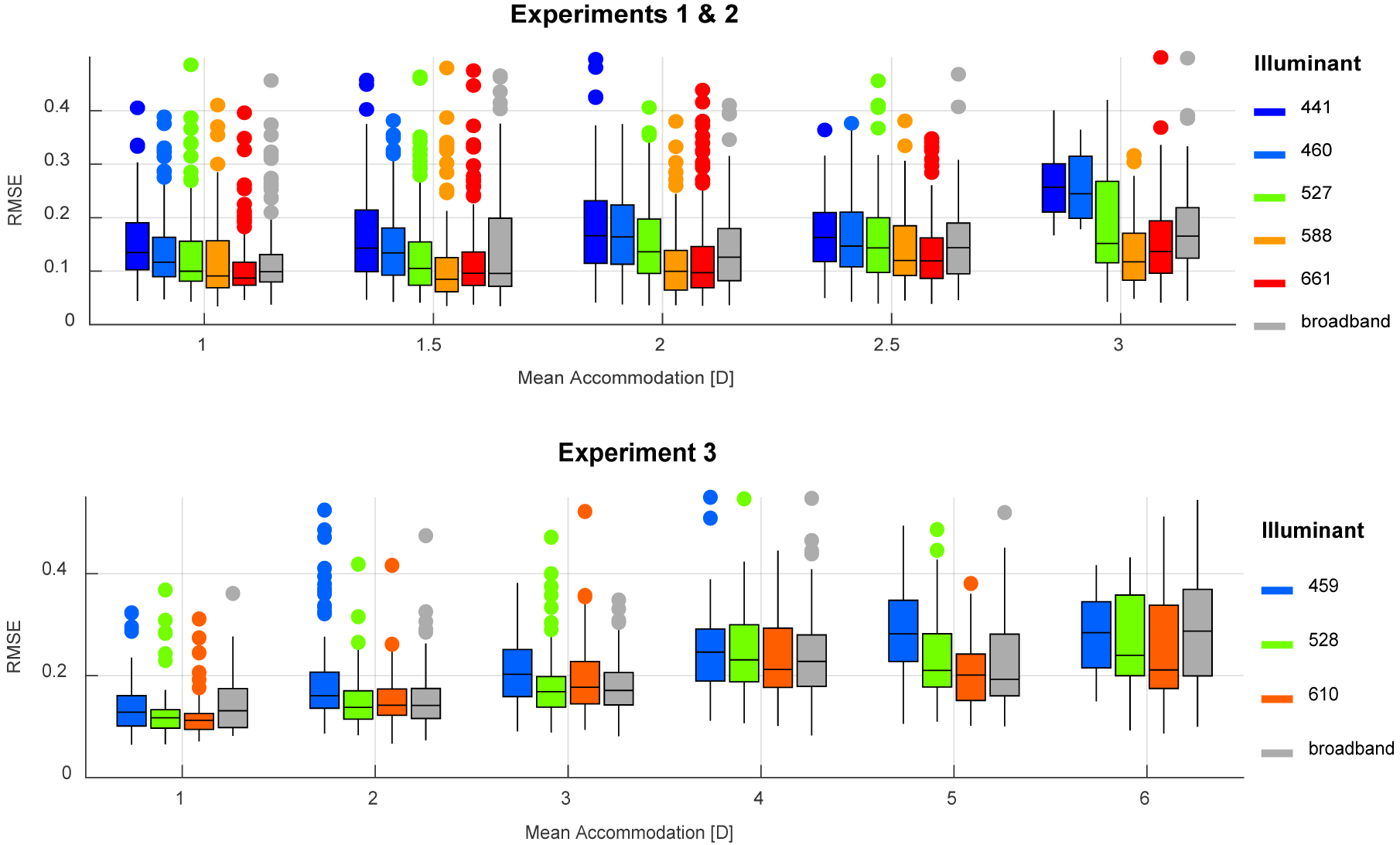
Distributions of the root-mean-square errors (RMSE) of an unconstrained linear fit through the within-trial accommodation response, as a function of mean accommodation and illuminant in experiments 1 and 2 (top) and experiment 3 (bottom). Each colour represents an illuminant as indicated by the legend. The mean accommodation values have been rounded and grouped for illustration purposes only. Note the wider range of the horizontal axis for Experiment 3. This figure can be generated using fig_h_RMSEplot.m in the code repository, and it uses the Gramm toolbox by Morel (2018).

To quantify these differences, we fitted two linear mixed effects models on the RMSEs, with mean accommodation as a continuous predictor and illuminant as a categorical predictor, while maintaining the full random-effects structure. The results are shown in the supplements, Supplementary Table 2.

In both experiments we observe similar intercepts of 0.15 dioptres (95% CI from 0.12 to 0.19) in experiments 1 and 2, and 0.13 dioptres (95% CI from 0.10 to 0.16) in experiment 3, with the 441nm and the 459nm illuminants, respectively. This means that at zero dioptres of refractive power, the accommodation response of observers to targets illuminated by these short wavelength illuminants fluctuates on average by 0.13 and 0.15 dioptres around the central response over time. The effect of refraction on RMSE was similar in both experiments as well, with one dioptre of increase in accommodation causing an increase of 0.02 dioptres in variability (95% CI from 0.01 to 0.03) in experiments 1 and 2, and an increase of 0.03 dioptres (95% CI from 0.02 to 0.04) in experiment 3. This agrees with several previous findings that the amplitude of accommodative microfluctuations increase with accommodation (Charman & Heron, 2015).

When comparing between different illuminants, we again see similar results in both datasets, with the highest within-trial variability in accommodation being observed for the shortest wavelength illuminants, and this variability decreasing as the peak wavelength of the illuminant increases. w

In summary, we see that the within-trial variability of the accommodation response increases with increasing refractive power, and it is lowest for the longer wavelength illuminants (588, 610 and 661nm) and highest for the shorter wavelength illuminants (441, 459 and 46w0nm). We found no systematic differences between broadband and narrowband illuminants, with the intra-trial variability being similar for the middle-wavelength (527 and 528nm) and broadband illuminants.

### Accommodation and pupil size

The median pupil diameter, centred to each participant, is illustrated as a function of accommodation and for the different illuminants used in Figure 12. To assess the effect of accommodation and of the different illuminants on the pupil diameter of participants, we fitted a linear mixed model for each experiment, with refraction in dioptres as a continuous predictor, and the illuminant as a categorical predictor, with random slopes and intercepts of participant. The latter were important as there was significant inter-individual variability in the median pupil diameter. The results are shown in Supplementary Table 3.

**Figure 12.**
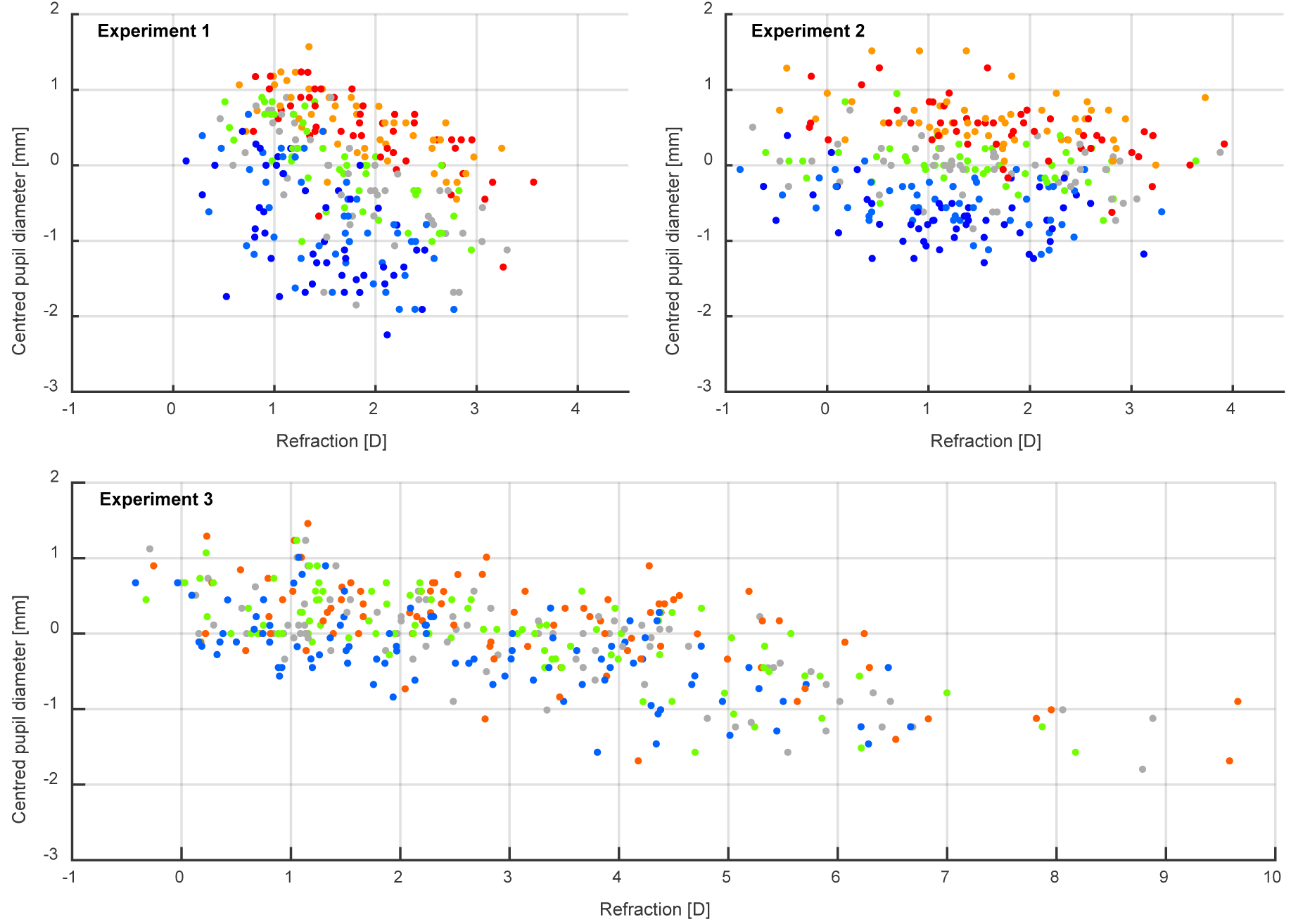
The median pupil diameter, centred to each participant, as a function of accommodation. For centring, we first subtract the mean pupil diameter, averaged over all trials recorded for that participant. We have plotted the median of this centred value, averaged over all trials for a given illuminant and participant. Each panel represents the data obtained in one experiment and the colour of the points represents the illuminant used. As illustrated, pupil diameter decreases more steeply with increasing accommodation in experiment 1 (where angular size was not kept constant) than in experiments 2 and 3. Note the wider range of the horizontal axis for Experiment 3. This figure can be generated with fig_e_PupilAcc.m in the code repository.

We found that pupil diameter significantly decreases as accommodation increases, although the rate of this change differs between experiments. In experiment 1 where the angular size of the stimuli increased as it was placed nearer the eye, pupil diameter decreased by 0.75mm (95% CI from 62 to 89mm) for every dioptre of increase in refraction. However, in experiments 2 and 3 where the angular size of the stimuli was kept constant, the slope was shallower, with pupil diameter decreasing by 0.16mm (95% CI from 0.06 to 0.26mm) and 0.18mm (95% CI from 0.06 to 0.29mm) for every 1D of increase in refraction, respectively.

The different illuminants used had a significant effect in the pupil size, with the shortest wavelength illuminants corresponding to the smallest diameters, and pupil size increasing progressively for longer wavelengths, even though the luminance was equal in all cases. The largest difference in pupil diameter for stimuli of equal luminance was of 1.70mm (95% CI: 1.38 to 2.01mm) for experiment 1 between the 441nm and 661nm illuminants, of 1.40mm (95% CI: 1.21 to 1.58mm) for experiment 2 between 441nm and 588nm, and of 0.45mm (95% CI: 0.30 to 0.59mm) for experiment 3 between 459nm and 610nm. Thus, a change in the peak wavelength of the illuminant used can have a larger effect on pupil diameter than changes in accommodation, particularly when the angular size of the stimuli is kept constant. Finally, the median pupil size for the broadband stimuli used seems to approximately correspond with the pupil diameter of middle wavelength illuminants: 527nm in experiments 1 and 2, and 528nm in experiment 3.

### Accommodative error and visual acuity

In experiment 3, the visual acuity of participants was measured for each illuminant at each distance, which allowed us to assess the effect that the median accommodation response of participants while they performed the staircase procedure had on their visual acuity. In Figure 13, we present the visual acuity thresholds obtained for all participants as a function of the median accommodative error (top) and as a function of the median pupil diameter (bottom). Individual figures for each participant are presented in the supplemental materials (Supplementary Figure 6).

**Figure 13.**
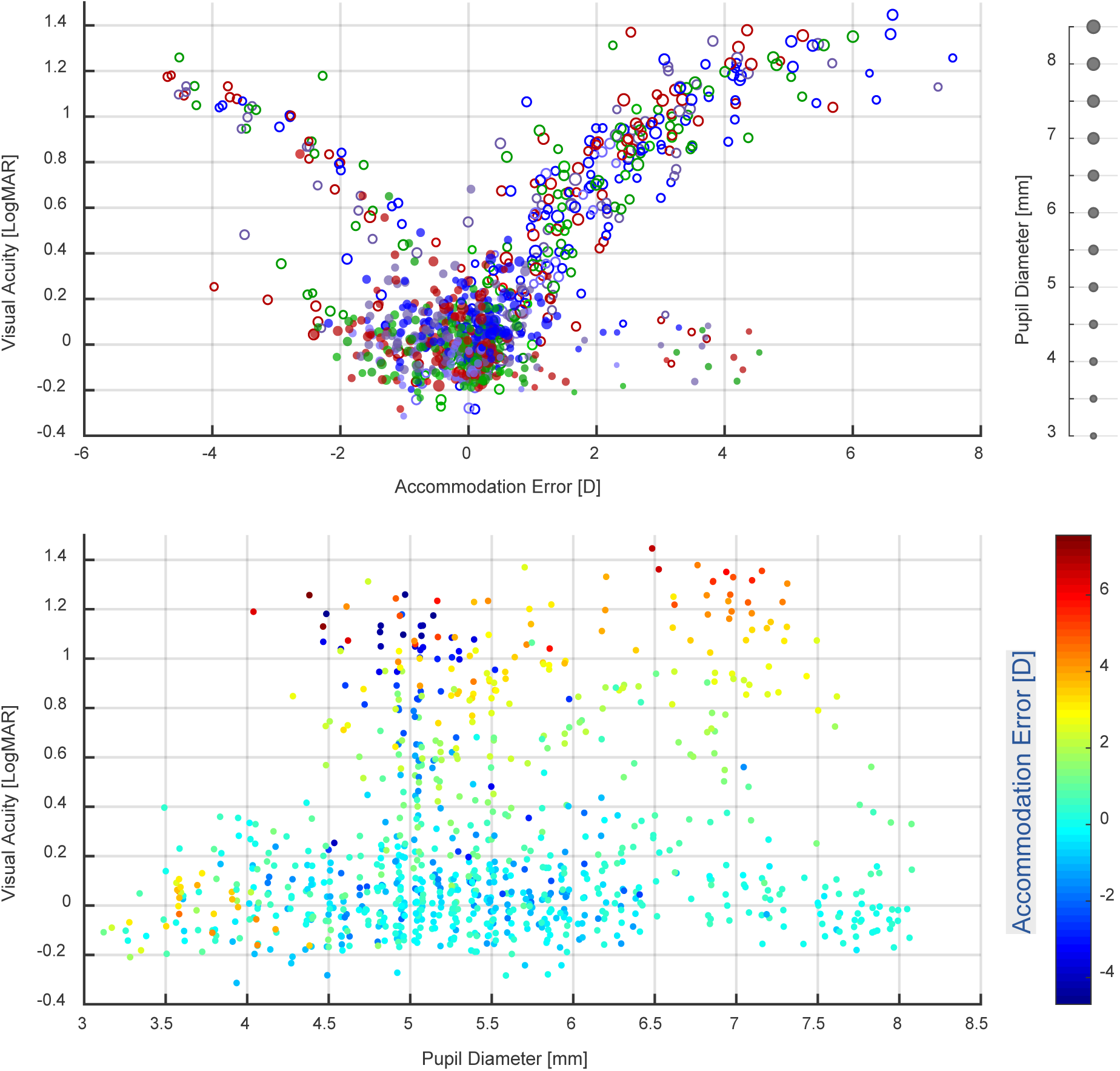
Visual acuity as a function of median accommodative error (top) and as a function of median pupil diameter (bottom). In the top panel, filled markers correspond to measurements over the linear portion of the accommodation response curve, and open markers measurements at distances where the function was saturated. The marker colours represent the illuminant used, and the marker sizes the corresponding median pupil size. In the bottom panel, the colour of the markers represents the median accommodative error. This figure can be generated with fig_g_VA_accError.m in the code repository.

Firstly, we analyse the results obtained over the linear portion of the accommodation response curve (see Figure 13 top, filled markers). As observed, over this portion, participants had visual acuity thresholds that were mostly concentrated between logMAR -0.2 and 0.2, which corresponds with better than normal to near normal vision. Median accommodative errors were mostly between -2D and 1D, with errors of larger magnitude mostly present for the portions where the response curve was found to be saturated. When looking at the individual results of each participant (see Supplementary Figure 6), we see that most of the data points with large negative errors of up to -2D and low visual acuity thresholds belong to subjects 16 and 19, which were the two participants that presented the typical lags in their accommodation response curves. Thus, it seems that in these two subjects, such lags did not correlate with a worsening of visual acuity.

Another relevant feature of the data is the small cluster of trials in which participants obtained low visual acuity thresholds between logMAR -0.2 and 0.2 despite presenting positive accommodative errors of up to 4.5D of magnitude. As illustrated in Figure 13 (bottom), one common feature of these trials is that the median pupil diameter of participants was mostly between 3 and 4mm. Smaller pupil sizes improve depth of focus which can decrease the effect that accommodative errors have on visual acuity. Additionally, the infrared photorefractor used relies on measuring the variation in reflected light intensity across the pupil to estimate the refractive state of the eye. This means that smaller pupils offer less information which could lead to less accuracy in the measurements taken. For these reasons, and since at these small pupil sizes the measured accommodative error does not seem to correlate with visual acuity thresholds, data points where the median pupil diameter was below 4mm were excluded from all analyses.

To further explore the relationship between accommodative error and visual acuity over the linear portion of the accommodation response curve, we fitted a linear mixed model with predictors of accommodative error magnitude, error sign, and their interaction, as well as illuminant, and random intercept and slopes of participant. The data used was the accommodative errors and visual acuity thresholds obtained within the accommodative range of participants (see filled markers in Figure 13), while excluding trials where the median pupil diameter was smaller than 4mm. The estimated coefficients are shown in Supplementary Table 4. We found that accommodative error magnitude was estimated to have a worsening effect on visual acuity, albeit the confidence intervals were wide, and the effect was not found to be significantly different from zero. The wide confidence intervals are likely reflecting the fact that the errors over the linear portion of the accommodation response curve were very small in magnitude for most subjects. In other words, subjects were accommodating successfully to the stimuli over a range of distances, resulting in small values of defocus and greater uncertainty in estimating its effect on visual acuity. However, the parameter estimates still indicate that the overall effect on visual acuity was detrimental, with thresholds worsening by LogMAR 0.10 for each dioptre of increase in negative accommodative error (95% CI from -0.05 to 0.24, *t*(8.37) = 1.51, *p* = 0.168), and by LogMAR 0.12 for each dioptre of increase in positive accommodative error (95% CI from 0.00 to 0.23, *t*(6.44) = 2.36, *p* = 0.053).

A significant effect of illuminant on visual acuity was found. When accommodative error is zero, visual acuity for the 459nm illuminant was estimated to be logMAR 0.04 (95% CI from -0.06 to 0.15). In comparison with this illuminant, visual acuity thresholds were lower for the 528nm illuminant by logMAR 0.11 (95% CI from 0.06 to 0.17, *t*(8.7) = 3.95, *p* = 0.004), for the 610nm illuminant by logMAR 0.08 (95% CI from 0.02 to 0.14, *t*(8.7) = 2.72, *p* = 0.024), and for the broadband illuminant by logMAR 0.09 (95% CI from 0.05 to 0.12, *t*(9.1) = 4.53, *p* = 0.001). Post-hoc pairwise comparisons of the estimated marginal means of visual acuity for each illuminant (i.e., the means averaged over the effects of accommodative error magnitude and sign), revealed that these differences were consistent and present across the small values of accommodative error found within the linear portion of the accommodation response curve. We found higher visual acuity thresholds for the 459nm illuminant when compared to the 528nmm illuminant by LogMAR 0.11 (95% CI from 0.02 to 0.20, *t*(8.91) = 3.87, *p* = 0.017), and by LogMAR 0.09 when compared to the broadband illuminant (95% CI from 0.02 to 0.15, *t*(8.74) = 4.34, *p* = 0.009). Visual acuity was also lower for the 610nm illuminant when compared to the 459nm one by LogMAR 0.08, although this difference was not significant (95% CI from -0.01 to 0.18, *t*(8.91) = 2.66, *p* = 0.100). This means that over the linear portion of the accommodation response curve and for equal values of accommodative error, visual acuity was worst for the shortest wavelength illuminant than for any of the other illuminants used. No significant differences were found in pairwise comparisons between the 610nm, 528nm and broadband illuminants.

To further explore the effect of accommodative error and the illuminants used we fitted linear mixed models on all the data obtained, that is, including distances that were nearer or farther away than participant’s accommodative range (see Figure 13, both open and filled markers). Due to the complexity of the data and the observed differences between the effect of underaccommodation (negative errors) and overaccommodation (positive errors), the dataset was separated accordingly and fitted separately. For positive accommodative errors, visual acuity thresholds seem to saturate for error magnitudes greater than 5.5D and at around logMAR 1.2; thus, these values (error magnitude > 5.5D and visual acuity > logMAR 1.2) were excluded from the analyses to improve model convergence. As with the previous model, trials where the median pupil diameter was less than 4mm were excluded, and the pupil diameter predictor was centred so that the intercept of the model was at 4mm. Several models were fitted to both datasets, with different combinations of accommodative error magnitude, pupil diameter, illuminant and retinal illuminance used as separate or interacting predictors, while maintaining the full structure of the random effects. Through multiple comparisons, it was determined that for both datasets, a model with predictors of error magnitude, pupil diameter, their interaction, and illuminant, had the greatest predictive power and lowest Akaike Information Criterion. The results of the fits for both datasets are shown in Supplementary Table 5.

For overaccommodation (Supplementary Table 5, top), we see that the accommodative error magnitude had a significant effect on visual acuity, with thresholds worsening by logMAR 0.21 for every 1D increase in error for a pupil diameter of 4mm (95% CI from 0.10 to 0.31, *t*(4.38) = 3.84, *p*=0.016). Furthermore, for every millimetre of pupil size increase, the effect of error magnitude on visual acuity significantly increases by LogMAR 0.08 (95% CI from 0.02 to 0.13, *t*(5.39) = 2.78, *p* = 0.036). This means that when participants have larger pupil sizes, their visual acuity is more affected as defocus increases. No significant differences in visual acuity were found between illuminants, so the differences previously observed for small accommodative errors within the linear portion of the accommodation response curve, are not present for positive accommodation errors of larger magnitude.

For underaccommodation (Supplementary Table 5s, bottom), we see that increases in error magnitude has a smaller effect on visual acuity that does not reach significance, with thresholds only worsening by logMAR 0.06 (95% CI from 0.00 to 0.12, *t*(9.97) = 1.82, *p* = 0.098) for every dioptre of increase in error and a pupil diameter of 4mm. For 1mm of increase in pupil diameter, the effect of accommodative error increases by logMAR 0.04 per dioptre, however, the confidence intervals are wide, and the effect is not significant (95% CI from -0.02 to 0.11, *t*(3.53) = 1.31, *p* = 0.268). Finally, visual acuity thresholds were higher for the 459nm when compared to the 528nm illuminant by LogMAR 0.07 (95% 0.01 to 0.14, *t*(7.31) = 2.16, *p*=0.066) and the broadband illuminant by 0.07 (95% CI 0.01 to 0.14, *t*(4.38) = 2.15, *p* = 0.086); however, these differences only reach significance at the 0.10 level.

The differences in results between both models could be explained by the fact that the negative accommodative errors were found mainly over the linear portion of the accommodation response curve, as the nearest distance used was not sufficient to reach the upper limit of the accommodative range of most participants. Indeed, we can see the similarities between the results for the linear portion of the accommodation response curve and for all the negative accommodative errors data. On the contrary, most participants did reach the lower limit of their accommodative range before the farthest distances used, so there was a wider range of data for the positive accommodative errors fit. However, it is possible that some of the differing results found are due to inherent differences in the effect of the accommodative error sign, as we see that for one of the two participants that reached their upper accommodative limit, visual acuity thresholds increased with a shallower slope when underaccommodating to the stimuli (see Supplementary Figure 6, Subject 2).

## Discussion

In this study we performed three experiments where we measured the steady-state accommodation and pupil responses of mostly untrained observers when looking at targets illuminated by different narrowband lights and placed at different distances. We found that most participants were able to accommodate under monochromatic light when the illumination of the target was changed abruptly and were able to maintain focus for the duration of the trials with similar accuracy as in white light, particularly at nearer distances. Like earlier workers, we found that the within-trial variability of the accommodation response increases with accommodation. We also found that variability increased slightly for shorter wavelengths at a given distance, perhaps reflecting the greater LCA (Figure 2). However, we found no systematic differences between broadband and narrowband illuminants in the variability of the accommodation response of observers over time, and the within-trial accommodative response fluctuated on average by similar amounts for the broadband and the green illuminants.

This finding contradicts some of the results reported by Kruger et al. (1997), as 38% of their sample had difficulty maintaining focus with narrowband targets placed away from the tonic state accommodation (at distances of 0 and 5 dioptres), while they could accommodate accurately to a broadband target at the same distance. These distances were included in our experiments, and we found no such impairment in steady-state accommodation. Instead, our results agree with those of (Atchison et al., 2004) that accommodative responses to targets with reduced spectral bandwidth were not more variable than responses to broadband targets; as well as with the finding of Charman & Tucker (1978) that participants can accommodate to narrowband stimuli of different wavelengths and maintain focus as accurately as in white light. It is notable that most of the participants in our sample were untrained naïve observers, as the one inexperienced observer of Charman & Tucker (1978) was not able to accommodate to the narrowband stimuli without additional training in the task. It is plausible that nowadays, with the increasing prevalence of narrowband LEDs as primaries in digital displays and as illumination sources, naïve observers have more experience accommodating to this type of stimulus and can make use of other cues to determine the sign and magnitude of the accommodative change, as well as to maintain focus. Some residual chromatic blur could still be present in our narrowband stimuli that could potentially serve as an accommodative cue since LEDs are not completely monochromatic (with a spectral bandwidth of ∼20nm). This would be especially true for shorter wavelengths, as the effects of LCA are greater towards the blue end of the spectrum. However, we did not find that accommodation was more accurate for short wavelengths, and in fact, the slope of the accommodation response curve as a function of distance was shallowest for these illuminants and the variability of the response higher. Thus, there is no evidence that any residual chromatic blur within a single narrowband illuminant contributed to the subject’s abilities to accommodate.

One of our main findings is that the slope of the linear portion of the accommodation stimulus-response curve becomes shallower as the peak wavelength of the narrowband illuminants decreases, which is caused by an increase in the difference in accommodation to different wavelengths as the target is placed at nearer distances. In other words, the extent to which participants change their accommodation responses to compensate for the LCA of the eye increases as they accommodate to nearer targets. At a distance of ∼0.5 dioptres there are no significant differences in the accommodation to different wavelengths, while at approximately 4.5 dioptres, participants change their accommodation responses nearly to the full extent that the chromatic eye model predicts (Thibos et al., 1992). This was a common finding in all three experiments, but there were considerable differences between participants, particularly at farther distances, as some subjects did change their accommodative responses to some extent to compensate for LCA even at 0.5 dioptres or farther. It is important not to confuse this with the nonlinear mapping from distance to dioptres.

Importantly, this does not just trivially reflect the nonlinear mapping from distance to dioptres. A range of ±0.1 dioptres centred on 0.5 dioptres (2m) corresponds to a distance of 83cm while the same range centred on 4.5 dioptres (22cm) corresponds to a distance of just 1cm. Thus it is natural for accommodation to be more precise *in metres* at near distances. However, as regards LCA, it is also more precise in dioptres.

Charman & Tucker (1978) found some comparable results. They measured the accommodation response curves for six participants under red and blue light and found that the slope was shallower for blue (468nm) in at least two of the subjects. However, for one of these subjects they measured the response to other narrowband illuminants (644nm, 579nm, 546nm, 503nm) and did not find a significant difference in the slope of the accommodation function other than for blue, albeit at distances of 1 and 2 dioptres this subject significantly underaccommodated for red (644nm). They theorized that this change could be partly explained by an increase in the LCA of the eye as the power of the crystalline lens increases. They then took objective measurements of the LCA of the eye in this participant and observed that it increased by ∼3% (0.03) per dioptre of accommodation, which they postulated could account for the results found for that subject (although some overaccommodation for blue and underaccommodation for red remained at the farthest distances tested of 1 and 2 dioptres, even after this adjustment). Across our sample, however, we found that the extent to which participants change their accommodation responses to correct for LCA increases by a much larger factor of 0.28 (95% CI from 0.08 to 0.45) per dioptre of increase in target vergence; thus, while an increase in LCA with accommodation might account for part of our results, it does not seem to fully explain them on its own. Charman & Tucker (1977) proposed that the change in slope in blue light might be due to reduced acuity at shorter wavelengths; however, we found that the difference in slope was significant between other illuminants tested as well (e.g., red and orange), so it does not seem to be unique to blue light.

Previous studies had found an increase in LCA with accommodation (Nutting & Larmor, 1997; Sivak & Millodot, 1974), as well as inter-individual differences in the LCA measured for different observers (Bedford & Wyszecki, 1957; Nutting & Larmor, 1997; Sivak & Millodot, 1974; Wald & Griffin, 1947). Sivak & Millodot (1974) in particular, used an achromatizing lens that corrected for most of the LCA of the eye (Bedford & Wyszecki, 1957), and subjectively measured the difference in optimal focal distance between different wavelengths at distances of 0.6, 3.0 and 7.1 dioptres. They observed that the difference in accommodation as a function of wavelength increased in all subjects from a mean of 0.40 dioptres at the farthest distance, to 0.65 dioptres at the nearest, with some variability between observers. If we perform a linear fit on their data, we see that the rate of increase in LCA is of 0.036 (or 3.6%) per dioptre of accommodation, similar to the results of Charman & Tucker (1977).

The fact that participants accommodate with increased accuracy to different wavelengths as the target nears is perhaps a surprising result, if we consider our finding that pupil size decreases with increasing accommodation, increasing the depth of focus of the eye. Indeed, it has been observed that the steady-state accommodative response of the eye is more accurate for larger pupil sizes (Ward & Charman, 1985), thus, we would expect observers to compensate for LCA to a greater extent when pupil size is larger at farther distances. In addition to this, in the first experiment the angular size of the target increased as it was placed nearer the observer, which would have decreased the high spatial frequency content of the image and increase power at lower spatial frequencies. Previous studies have found that the steady-state accommodation response is more accurate for higher spatial frequencies and substantial in error for lower spatial frequencies (Charman & Tucker, 1977), so we would expect this factor to contribute to responses being less accurate at nearer distances, but the results of this experiment indicate otherwise.

One possibility that could explain these results is that, as LCA is significantly reduced in narrowband light, participants are making use of other cues to find the optimal focal distance for different wavelengths, and these cues might change with target vergence. Specifically, the micro-fluctuations of the crystalline lens have been found to increase in magnitude as accommodation increases due to the increased freedom of movement (Day et al., 2006; Kotulak & Schor, 1986; Stark & Atchison, 1997), covering an approximate range of 0.02 dioptres in both directions when the mean accommodation is 1 dioptre, and increasing to a range of up to 0.1 dioptres when the accommodative response is 4 dioptres. These micro-fluctuations could serve as a cue to accommodation by providing negative feedback to the accommodative control mechanism, essentially functioning as sub-threshold blur detector; thus, it is possible that the increased range of these micro-fluctuations at higher accommodation levels allows the visual system to find the focal distance for each wavelength more accurately when the colour of the target is changed and in the absence of the chromatic blur caused by LCA. However, this is only speculation on our part, as there is no evidence in the literature that the increased amplitude of micro-fluctuations can lead to higher accommodative accuracy, and on the contrary, consistent steady-state errors when accommodating to nearer targets are often found (Plainis et al., 2005).

Another factor that could be influencing these results is that accommodation might become less accurate as observers reach the far point of their accommodative range, which would happen at nearer physical distances for targets illuminated by shorter wavelength light. In other words, the typical lag of the accommodation response curve would start to occur at nearer distances for short than for long wavelength light, which would cause the slope estimate to be shallower. However, the analysis performed to determine the accommodative range of each participant for each individual illuminant, and the subsequent exclusion of the saturated points should have addressed this issue, at least partially. Overall, our results seem novel within the literature, although Charman & Tucker (1978) had some comparable findings with two of their subjects. It is not clear why participants increasingly correct for LCA at nearer distances when accommodating to narrowband stimuli, and more research is needed in this area to explain these results, as well as to further explore the individual differences between observers.

Another of our findings was that accommodation to white light tended to overlap with middle wavelengths over all distances tested (see Figure 5 to Figure 8). When the targets were illuminated by a white light with the highest luminous spectral power between 530 and 590nm, accommodation was similar to the narrowband illuminants of similar peak wavelengths (527 and 588nm) over all distances tested, although the slope as a function of distance was steeper than for the narrowband illuminants and closer to one, with accommodation slightly shifting from green towards orange as target vergence increased from 0.5 to 3 dioptres. In a third experiment where a broadband illuminant was created by using the three narrowband primaries at equal luminance, accommodation seemed to overlap with the red illuminant (610nm), or between the red and green (528nm) illuminants, over most distances tested.

Some previous studies have investigated the wavelength that comes into focus in the retina in broadband white light at different distances. Ivanoff (1949) found that with increasing accommodation, the wavelength that was kept in focus in the retina decreased, from ∼600nm at 0.5 dioptres, to ∼500nm at 2.5 dioptres. Similarly, Sivak & Millodot (1974) found that the wavelength in focus changed from 620nm at a distance of 0.7 dioptres, to 530nm at a distance of 7.1 dioptres. Ivanoff (1949) proposed that this change of wavelength-in-focus with distance could explain the lag and leads of the accommodative response by a process of “sparing of accommodation”, that is, the visual system uses the LCA of the eye to accommodate as close as possible to the tonic or resting state, choosing to accommodate to shorter wavelengths at near distances, as they require the least refractive power, and to longer wavelengths for farther distances. If this were the case, one would expect to find much steeper stimulus-response curves for narrowband illuminants than for white light, which does not agree with our findings. Similarly, Charman & Tucker (1978) and Jaskulski et al. (2016) did not find that the stimulus-response curves for narrowband light of different wavelengths was steeper than for white light. Thus, no “sparing of accommodation” seems to be taking place, and it is possible that those earlier findings were due to the spherical aberration of the eye usually changing from positive at far, to increasing negative values as accommodation increases (Del Águila-Carrasco et al., 2020; Thibos et al., 2013). When spherical aberration is positive at far distances, the rays entering through the periphery of the pupil will come into focus in front of the retina, so the shorter wavelength content of that light will be more out of focus and longer wavelengths will come into focus closer to the retina. When accommodation increases and spherical aberration becomes negative, the peripheral rays will come into focus behind the retina, so the longer wavelength content will be more out of focus and shorter wavelengths more in focus. Thus, it is possible that the phenomenon observed by Ivanoff (1949) was due to the distribution of light in the retina changing due to a change in the sign of the spherical aberration of the eye, rather than by the visual system shifting the wavelength that is kept in focus in white light.

Our results seem to indicate that when accommodating to white light, the wavelength that is kept in focus is between 527 and 610nm, which agrees with findings by DeHoog & Schwiegerling (2007) that the best focus in white light corresponded to best focus for monochromatic light between 590 and 610nm. The slightly steeper slope, closer to unity, that we found for white light when compared to the green or orange illuminants could be because chromatic blur due to LCA aids accommodation.

Another of our findings was that steady-state median pupil diameter decreases with increasing accommodative state and with decreasing peak wavelength in narrowband illuminants, even when luminance and angular size was equal (see Figure 12). The effect of wavelength on narrowband illuminants can be explained by the contribution of the melanopsin photopigment present in some retinal ganglion cells (ipRGCs) to steady-state pupil size control (Spitschan, 2019b, 2019a). While our stimuli were created to provide equal input to the luminance channel, pupil size control has a strong input from the ipRGCs in addition to the cone photoreceptors. The melanopsin photopigment is more sensitive to short wavelength light than the L and M cones, with a peak sensitivity at 480nm (Enezi et al., 2011); thus, the shorter wavelength light used in our experiments would provide greater stimulation to the ipRGCs and the pupil control mechanism than the longer wavelength illuminants of equal luminance.

The literature investigating the effect of accommodation on pupil size offers a less clear picture to explain our results. While the near triad of accommodation, convergence and pupil constriction is a well-established fact, there is contradictory evidence on whether convergence or accommodation are responsible for the pupil response at near distances. Some studies have found that accommodation alone does not trigger a pupil response when convergence and other factors such as target size and alignment are controlled (Feil et al., 2017; Phillips et al., 1992; Stakenburg, 1991). However, one of these studies measured dynamic rather than steady-state pupil responses, and another did not directly measure accommodation, but inferred it from acuity measurements. Other studies have arrived to the opposite conclusion, finding that blur-driven accommodation and not fusional vergence, cause pupil constriction (Marg & Morgan, 1949; Phillips et al., 1992; Wilson, 1973). As to the extent of the change, Marg & Morgan (1950) found that pupil diameter changed on average by 0.48mm per dioptre of accommodative stimulus and did not change with convergence, although it is possible that factors such awareness of target proximity might have played a role (Phillips et al., 1992). On the other hand, O’Neill & Stark (1968) and van der Wildt & Bouman (1971) both reported measurements showing an increase of ∼0.17mm per dioptre of accommodation when describing the design and construction of equipment to measure accommodation, vergence and pupil diameter dynamically. While their experiments had more carefully controlled parameters, presenting the targets monocularly and maintaining constant target size, alignment along the axis of the eye, and luminance, their sample was limited to just one subject each. Here, we present results with a larger sample that show similar estimates, with steady-state pupil diameter decreasing by 0.16 to 0.18mm per dioptre of accommodation for targets that were viewed monocularly, aligned with the axis of the stimulated eye, and with constant luminance and angular size. Furthermore, in one of our experiments the apparent distance of the target was also kept constant, with the accommodation being driven by placing lenses in front of the eye. Thus, our results provide further support to the idea that steady-state pupil constriction can be caused by accommodation alone, although the rate of change is smaller than reported in some of the previous studies.

Over the quasi-linear portion of the accommodation response curve, we found that accommodative errors (i.e., the difference between accommodative demand and the median response) had mostly magnitudes of up to 1 dioptre in either direction, although underaccommodation was more prevalent in our sample. An interesting finding was that not all subjects presented the consistent lags in accommodation as the target nears that are often reported in the literature (Nakatsuka et al., 2003), and overall, there was great inter-subject variability in the shape and slope of the stimulus-response curve. Furthermore, the two subjects that did present significant lags of up to 1.5 and 2 dioptres of magnitude at near distances in the third experiment, did not have their visual acuity significantly impaired by those accommodative errors. In fact, over the linear portion of the accommodation response curve for all participants, we found that accommodative error did not have a significant effect on visual acuity thresholds.

It is possible that our measurements were not precise enough to capture the relationship between accommodative error and visual acuity for a relatively small range of errors. Accommodation was measured as participants performed the staircase procedure with targets of different spatial frequency being presented and pupil size allowed to change freely, so the accommodative response would not be the only factor affecting retinal image quality, and the median of this response might not be representative of the defocus of the retinal image when participants were viewing the smaller targets that were more critical to the thresholds obtained. The depth of field of the eye would also allow some of these accommodative errors to not have a detrimental effect on visual acuity. For pupil diameters between 4 and 6mm, the depth of field can be between 0.4 and 0.5 dioptres, even for high spatial frequencies and monochromatic light (Marcos et al., 1999), albeit higher estimates have been obtained (Wang & Ciuffreda, 2006). Jaskulski et al. (2016), for example, found that the subjective depth of field for a pupil diameter of 3.8mm and a target of LogMAR 0.1 size, was approximately 1.19 dioptres for narrowband light, and slightly higher for polychromatic light. Of course, even the higher estimates are not enough to fully explain the results obtained, particularly in the two subjects that showed lags of significant magnitude.

Recently, Labhishetty et al. (2021) used several objective and subjective measurements to measure the accommodation of the eye. They found that, for target distances between 0 and 6 dioptres, objective measurements had higher accommodative errors than subjective measurements based on visual acuity. In particular, the measurements taken using a photorefractor gave the largest measured lags, with magnitudes between 0.5 and 1.5 dioptres. Despite these large errors, subjective measurements indicated that participants were accommodating accurately to the distance that maximised their visual acuity, with the subjective errors being much lower at ∼0.15 dioptres. These results are comparable to ours, as we used a photorefractor to measure accommodation and found errors of considerable magnitude (mostly lags) that did not seem to have a detrimental effect on visual acuity. In particular, the two subjects that displayed the more typical large accommodative lags, maintained visual acuity thresholds close to logMAR 0 regardless of the magnitude of these errors.

It has been suggested that the consistent errors that are observed when accommodation is measured objectively with a photorefractor (i.e., lags and leads), might actually be the consequence of the spherical aberration of the eye, particularly its change in sign with accommodation (Plainis et al., 2005; Thibos et al., 2013). As mentioned previously, the eyes of most observers tend to exhibit positive spherical aberration when accommodating at far distances, which decreases steadily with increasing demand and becomes negative at nearer distances (Del Águila-Carrasco et al., 2020). This means that peripheral rays will come into focus in front of the retina at far distances and behind the retina at near. As photorefractors use the distribution of reflected light across the entire pupil to estimate the refractive state of the eye, they might put more weight on these marginal rays than the visual system does, leading to apparent leads and lags in accommodation, even when paraxial rays are focused correctly in the retina (Thibos et al., 2013). Thus, it is possible that the large accommodative lags observed in two of the subjects in our third experiment are due to their own spherical aberration and the method used to measure accommodation.

While accommodative errors were small over the linear portion of the accommodation response curve, and thus, did not have a significant effect on visual acuity, we did find differences caused by the illuminants used. Visual acuity thresholds were significantly lower for blue light when compared to the other three illuminants. When accommodative error was zero, the visual acuity for blue light was estimated to be LogMAR 0.04, while for the red, green, and broadband illuminants it was LogMAR -0.04, -0.07 and -0.05, respectively. As luminance was kept the same for all illuminants, giving the same input to the luminance (L+M) channel, the differences in visual acuity cannot be explained by the differences in sensitivity to different wavelengths. The lower visual acuity for blue light can rather be explained by the blur caused by LCA for shorter wavelengths. For a spectral distribution that is not completely monochromatic, the LCA of the eye will cause greater defocus at shorter wavelengths, which will in turn reduce retinal image contrast, particularly at higher spatial frequencies. The blue light used had a spectral bandwidth (full width at half maximum) of 20nm around a peak wavelength of 459nm, which would cause a difference in defocus of 0.21 dioptres. In comparison, even though the green and red illuminants had slightly larger spectral bandwidths (25 and 28nm, respectively), there would only be a difference of 0.16 and 0.11 dioptres in defocus across the bandwidth of their respective spectral distributions. As this defocus is caused by the intrinsic change in refractive index of the eye with wavelength, the accommodative response alone cannot correct it. Furthermore, it seems that the smaller average pupil size for blue light is not sufficient to improve focus or the retinal contrast of the image. Indeed, it has been observed previously that, while visual acuity in blue narrowband light is lower under normal conditions, compensating for the LCA of the eye improves the thresholds so that they more closely match those obtained in green, red and white light (Domenech et al., 1994). Overall, however, our results indicate that the visual acuity thresholds for blue were still within the range of normal vision, and the differences with the other primaries of the display were small (∼LogMAR 0.09), such that it is unlikely to have a significant impact in most real-life applications.

The increased defocus caused by LCA for shorter wavelengths could also explain the increased variability of the accommodation response for these illuminants when compared to longer wavelength ones. It has been previously reported that the magnitude of the micro-fluctuations of accommodation increases with increasing blur in the image (Niwa & Tokoro, 1998) and with decreasing contrast (Charman & Heron, 2015), and correlate with the objective depth-of-focus of the eye (Yao et al., 2010). The higher defocus caused for narrowband LEDs of shorter peak wavelength would reduce retinal image contrast and increase the depth-of-focus, which could increase the magnitude of the micro-fluctuations of accommodation, resulting in the higher within-trial variability of the response observed. We did not find, however, an increased variability in the broadband illuminants used, even though defocus and depth-of-focus would be greater in this condition (Jaskulski et al., 2016). Niwa & Tokoro (1998) previously found that micro-fluctuations increase as the blur of the image increases, but only for small amounts of blur. As the magnitude of blur continues to increase, the magnitude of the micro-fluctuations starts to decrease again, and the amount of blur at which micro-fluctuations peak is lower for higher spatial frequencies. These changes were found in particular for the low-frequency components of micro-fluctuations, which are caused by the action of the ciliary muscles on the crystalline lens (i.e., are under neural control), and may be of more significance to accommodative control (Charman & Heron, 2015). As blur increases, micro-fluctuations might increase in order to serve as an error signal to accommodation and improve the accuracy of the response. With higher depth-of-focus, the magnitude of the micro-fluctuations would need to be higher in order to provide the same amount of information for error detection to the accommodative control mechanism. (Jaskulski et al., 2016). Niwa & Tokoro (1998) postulate that when blur surpasses a certain threshold, it can no longer be discriminated and there is an overall reduction in micro-fluctuations, with this threshold being lower for higher spatial frequencies as they are more affected by defocus. Thus, it is possible that the higher defocus caused by LCA for the broadband illuminants is not detectable, causing the magnitude of the micro-fluctuations to be lower when compared to the short wavelength illuminants. It is important to note that these differences in response variability were found even after controlling for the mean accommodative state of the eye, as micro-fluctuations have been found to increase with increasing accommodation (Day et al., 2006; Kotulak & Schor, 1986; Stark & Atchison, 1997), which we corroborated in this study.

Interestingly, visual acuity thresholds in white light did not differ significantly from those obtained with the narrowband green and red illuminants. The chromatic blur caused by LCA in broadband white light did not seem to impair visual acuity, even though for our broadband illuminant all three primaries were set at equal luminance (so the chromatic blur would not be attenuated by the reduced luminous sensitivity at longer and shorter wavelengths). Domenech et al. (1994) found comparable results: with normal LCA, the visual acuity in white was similar as for red and green narrowband light and compensating for the LCA of the eye did not significantly improve the thresholds for white light. More recently, Suchkov et al. (2019) also found that correcting the LCA of the eye did not cause the predicted improvement in visual acuity, but rather a slight decrease (albeit not statistically significant), and that doubling the LCA of the eye had a more detrimental effect than predicted from their simulations. Further evidence from these authors also shows that correcting for the LCA of the eye does not improve visual acuity in high contrast conditions, even when subjects are given time to adapt to the corrected LCA (Fernandez et al., 2020). Thus, it seems that the ability of the visual system to resolve small targets with precision is not impaired by the chromatic blur caused by LCA, at least in high contrast conditions.

Finally, when considering all the visual acuity measurements, including those obtained beyond the accommodative range of participants, we found that overaccommodation had a significant detrimental effect on visual acuity, and a statistically significant interaction with pupil diameter, such that visual acuity was more affected by defocus in larger pupils than in smaller ones. This is consistent with previous findings in the literature of an increased depth of focus with smaller pupil sizes (Marcos et al., 1999; Wang & Ciuffreda, 2006). No effect of illuminant was found, indicating that larger values of defocus affect broadband and narrowband targets equally, including blue light. This is also consistent with previous findings that indicate that depth of focus increases with decreasing acuity (Tucker & Charman, 1975), such that the small amount of blur caused by LCA for blue light would no longer have an impact on retinal image quality.

In summary, we found that narrowband illumination can be an adequate stimulus to accommodation when compared to white light, even in a sample of mostly untrained observers. We also found that the extent to which participants change their accommodative responses to compensate for the LCA of the eye increases at nearer distances and matches the predictions of the chromatic eye model from approximately 4.5 dioptres (22cm) and nearer; a finding which is not fully explained by the previously reported increase in LCA of ∼3% per dioptre of accommodation. This means that considering the spectral distribution of the display primaries and its effect on accommodation might be more relevant for displays that are used at nearer distances, such as mobile phones or computer monitors, than for those that are viewed farther away such as television or cinema screens. We found no detrimental effects on visual acuity for narrowband light, with only blue light causing a significant worsening of the thresholds due to the larger spread of defocus caused by LCA at shorter wavelengths for a display primary that is not completely monochromatic. However, visual acuity in blue light was still within the range of normal vision (∼LogMAR 0), and this small difference is unlikely to be relevant to real-life display applications where images with multiple spatial frequencies are used.

## Data and code availability

Data are available at https://figshare.com/s/71b68d133237332b9d9f, and Matlab analysis code is available at https://figshare.com/s/a310d6bc0415e4f55d10.

## Acknowledgements

This research was funded by the European Union’s Horizon 2020 research and innovation program under the Marie Skłodowska-Curie grant agreement No. 676401, European Training Network on Full Parallax Imaging. Abigail Finch acknowledges a PhD studentship funded by the Engineering and Physical Sciences Research Council (EPSRC). For the purpose of Open Access, the authors have applied a CC BY public copyright licence to any Author Accepted Manuscript (AAM) version arising from this submission.

## Author contributions (CREDIT taxonomy)

|  |  |
| --- | --- |
| Conceptualization | MFA, APF, GL, JCAR |
| Data curation | MFA |
| Formal Analysis | MFA |
| Funding acquisition | GL, JCAR |
| Investigation | MFA, APF |
| Methodology | MFA, APF |
| Project administration | GL, JCAR |
| Resources |  |
| Software | MFA, APF |
| Supervision | GL, JCAR |
| Validation | MFA, APF |
| Visualization | MFA, APF |
| Writing – original draft | MFA |
| Writing – review & editing | MFA, GL, JCAR |

## References

Aggarwala, K. R., Kruger, E. S., Mathews, S., & Kruger, P. B. (1995). Spectral bandwidth and ocular accommodation. JOSA A, 12(3), 450–455. https://doi.org/10.1364/JOSAA.12.000450

Aggarwala, K. R., Nowbotsing, S., & Kruger, P. B. (1995). Accommodation to monochromatic and white-light targets. Investigative Ophthalmology & Visual Science, 36(13), 2695–2705.

Atchison, D. A., Strang, N. C., & Stark, L. R. (2004). Dynamic accommodation responses to stationary colored targets. Optometry and Vision Science.

Bates, D., Maechler, M., Bolker [aut, B., cre, Walker, S., Christensen, R. H. B., Singmann, H., Dai, B., Scheipl, F., Grothendieck, G., Green, P., Fox, J., Bauer, A., & simulate.formula), P. N. K. (shared copyright on. (2023). lme4: Linear Mixed-Effects Models using “Eigen” and S4 (1.1-33). https://cran.r-project.org/web/packages/lme4/index.html

Bedford, R. E., & Wyszecki, G. (1957). Axial Chromatic Aberration of the Human Eye. JOSA, 47(6), 564_1-565. https://doi.org/10.1364/JOSA.47.0564_1

Bharadwaj, S. R., Sravani, N. G., Little, J.-A., Narasaiah, A., Wong, V., Woodburn, R., & Candy, T. R. (2013). Empirical variability in the calibration of slope-based eccentric photorefraction. Journal of the Optical Society of America. A, Optics, Image Science, and Vision, 30(5), 923–931.

Bobier, W. R., Campbell, M. C., & Hinch, M. (1992). The influence of chromatic aberration on the static accommodative response. Vision Research, 32(5), 823–832. https://doi.org/10.1016/0042-6989(92)90025-e

Campbell, R. (2023). Raacampbell/shadedErrorBar [Matlab]. https://github.com/raacampbell/shadedErrorBar

Charman, W. N. (1989). Accommodation performance for chromatic displays. Ophthalmic & Physiological Optics: The Journal of the British College of Ophthalmic Opticians (Optometrists), 9(4), 459–463.

Charman, W. N., & Heron, G. (2015). Microfluctuations in accommodation: An update on their characteristics and possible role. Ophthalmic and Physiological Optics, 35(5), 476–499. https://doi.org/10.1111/opo.12234

Charman, W. N., & Tucker, J. (1977). Dependence of accommodation response on the spatial frequency spectrum of the observed object. Vision Research, 17(1), 129–139. https://doi.org/10.1016/0042-6989(77)90211-5

Charman, W. N., & Tucker, J. (1978). Accommodation and color. Journal of the Optical Society of America, 68(4), 459–471. https://doi.org/10.1364/josa.68.000459

Day, M., Strang, N. C., Gray, L. S., & Seidel, D. (2006). Depth of Focus, Refractive Error and Accommodation Microfluctuations. Investigative Ophthalmology & Visual Science, 47(13), 3892–3892.

DeHoog, E., & Schwiegerling, J. (2007). Position of White Light Best Focus in the Human Eye. Investigative Ophthalmology & Visual Science, 48(13), 993.

Del Águila-Carrasco, A. J., Kruger, P. B., Lara, F., & López-Gil, N. (2020). Aberrations and accommodation. Clinical and Experimental Optometry, 103(1), 95–103. https://doi.org/10.1111/cxo.12938

Domenech, B., Seguí, M. M., Capilla, P., & Illueca, C. (1994). Variation of the visual acuity—Luminance function with background colour. Ophthalmic & Physiological Optics: The Journal of the British College of Ophthalmic Opticians (Optometrists), 14(3), 302–305. https://doi.org/10.1111/j.1475-1313.1994.tb00013.x

Donohoo, D. T. (1985). Accommodation With Displays Having Color Contrast [PhD Thesis]. Virginia Polytechnic Institute and State University.

Enezi, J. A., Revell, V., Brown, T., Wynne, J., Schlangen, L., & Lucas, R. (2011). A “melanopic” spectral efficiency function predicts the sensitivity of melanopsin photoreceptors to polychromatic lights. Journal of Biological Rhythms, 26(4), 314–323. https://doi.org/10.1177/0748730411409719

Feil, M., Moser, B., & Abegg, M. (2017). The interaction of pupil response with the vergence system. Graefe’s Archive for Clinical and Experimental Ophthalmology = Albrecht Von Graefes Archiv Fur Klinische Und Experimentelle Ophthalmologie, 255(11), 2247–2253. https://doi.org/10.1007/s00417-017-3770-2

Fernandez, E. J., Suchkov, N., & Artal, P. (2020). Adaptation to the eye’s chromatic aberration measured with an adaptive optics visual simulator. Optics Express, 28(25), 37450–37458. https://doi.org/10.1364/OE.404296

Fincham, E. F. (1951). The Accommodation Reflex and its Stimulus. The British Journal of Ophthalmology, 35(7), 381–393.

Ghahghaei, S., Reed, O., Candy, T. R., & Chandna, A. (2019). Calibration of the PlusOptix PowerRef 3 with change in viewing distance, adult age and refractive error. Ophthalmic and Physiological Optics, 39(4), 253–259. https://doi.org/10.1111/opo.12631

Ivanoff, A. (1949). Focusing Wave-Length for White Light. JOSA, 39(8), 718–718. https://doi.org/10.1364/JOSA.39.000718

Jaskulski, M., Marín-Franch, I., Bernal-Molina, P., & López-Gil, N. (2016). The effect of longitudinal chromatic aberration on the lag of accommodation and depth of field. Ophthalmic and Physiological Optics, 36(6), 657–663. https://doi.org/10.1111/opo.12320

Kingdom, F., & Prins, N. (2016). Psychophysics—2nd Edition. Academic Press. https://www.elsevier.com/books/psychophysics/kingdom/978-0-12-407156-8

Kleiner, M., Brainard, D., Pelli, D., Ingling, A., Murray, R., & Broussard, C. (2007). What’s new in Psychtoolbox-3. Perception, 36(14), 1.

Kotulak, J. C., Morse, S. E., & Billock, V. A. (1995). Red-green opponent channel mediation of control of human ocular accommodation. The Journal of Physiology, 482 (Pt 3)(Pt 3), 697–703. https://doi.org/10.1113/jphysiol.1995.sp020552

Kotulak, J. C., & Schor, C. M. (1986). Temporal variations in accommodation during steady-state conditions. Journal of the Optical Society of America. A, Optics and Image Science, 3(2), 223–227. https://doi.org/10.1364/josaa.3.000223

Krinov, E. L. (1953). Spectral Reflectance Properties of Natural Formations. National Research Council of Canada. https://books.google.co.uk/books/about/Spectral_Reflectance_Properties_of_Natur.html?id=TtivnQEACAAJ&redir_esc=y

Kruger, P. B., Aggarwala, K. R., Bean, S., & Mathews, S. (1997). Accommodation to stationary and moving targets. Optometry and Vision Science: Official Publication of the American Academy of Optometry, 74(7), 505–510. https://doi.org/10.1097/00006324-199707000-00018

Kruger, P. B., Mathews, S., Aggarwala, K. R., & Sanchez, N. (1993). Chromatic aberration and ocular focus: Fincham revisited. Vision Research, 33(10), 1397–1411. https://doi.org/10.1016/0042-6989(93)90046-y

Labhishetty, V., Cholewiak, S. A., Roorda, A., & Banks, M. S. (2021). Lags and leads of accommodation in humans: Fact or fiction? Journal of Vision, 21(3), 21–21. https://doi.org/10.1167/jov.21.3.21

Lenth, R. V., Bolker, B., Buerkner, P., Giné-Vázquez, I., Herve, M., Jung, M., Love, J., Miguez, F., Riebl, H., & Singmann, H. (2023). emmeans: Estimated Marginal Means, aka Least-Squares Means (1.8.6). https://cran.r-project.org/web/packages/emmeans/index.html

Lovasik, J. V., & Kergoat, H. (1988a). Accommodative performance for chromatic displays. Ophthalmic & Physiological Optics: The Journal of the British College of Ophthalmic Opticians (Optometrists), 8(4), 443–449. https://doi.org/10.1111/j.1475-1313.1988.tb01183.x

Lovasik, J. V., & Kergoat, H. (1988b). The effect of optical defocus on the accommodative accuracy for chromatic displays. Ophthalmic & Physiological Optics: The Journal of the British College of Ophthalmic Opticians (Optometrists), 8(4), 450–457. https://doi.org/10.1111/j.1475-1313.1988.tb01184.x

MacKenzie, K. J., Hoffman, D. M., & Watt, S. J. (2010). Accommodation to multiple-focal-plane displays: Implications for improving stereoscopic displays and for accommodation control. J Vis, 10(8), 22. https://doi.org/10.8.22 [pii] 10.1167/10.8.22

Marcos, S., Moreno, E., & Navarro, R. (1999). The depth-of-field of the human eye from objective and subjective measurements. Vision Res, 39(12), 2039–2049.

Marg, E., & Morgan, M. W. (1949). The pupillary near reflex; the relation of pupillary diameter to accommodation and the various components of convergence. American Journal of Optometry and Archives of American Academy of Optometry, 26(5), 183–198.

Marg, E., & Morgan, M. W. (1950). Further investigation of the pupillary near reflex; the effect of accommodation, fusional convergence and the proximity factor on pupillary diameter. American Journal of Optometry and Archives of American Academy of Optometry, 27(5), 217– 225.

Morel, P. (2018). Gramm: Grammar of graphics plotting in Matlab. Journal of Open Source Software, 3(23), 568. https://doi.org/10.21105/joss.00568

Nakatsuka, C., Hasebe, S., Nonaka, F., & Ohtsuki, H. (2003). Accommodative lag under habitual seeing conditions: Comparison between adult myopes and emmetropes. Japanese Journal of Ophthalmology, 47(3), 291–298. https://doi.org/10.1016/s0021-5155(03)00013-3

Niwa, K., & Tokoro, T. (1998). Influence of spatial distribution with blur on fluctuations in accommodation. Optometry and Vision Science: Official Publication of the American Academy of Optometry, 75(3), 227–232. https://doi.org/10.1097/00006324-199803000-00029

Nutting, P. G., & Larmor, J. (1997). The axial chromatic aberration of the human eye. *Proceedings of the Royal Society of London. Series A*, Containing Papers of a Mathematical and Physical Character, 90(620), 440–443. https://doi.org/10.1098/rspa.1914.0070

O’Neill, W. D., & Stark, L. (1968). Triple-function ocular monitor. Journal of the Optical Society of America, 58(4), 570–573. https://doi.org/10.1364/josa.58.000570

Phillips, N. J., Winn, B., & Gilmartin, B. (1992). Absence of pupil response to blur-driven accommodation. Vision Research, 32(9), 1775–1779. https://doi.org/10.1016/0042-6989(92)90170-n

Plainis, S., Ginis, H. S., & Pallikaris, A. (2005). The effect of ocular aberrations on steady-state errors of accommodative response. Journal of Vision, 5(5), 7–7. https://doi.org/10.1167/5.5.7

Seidemann, A., & Schaeffel, F. (2002). Effects of longitudinal chromatic aberration on accommodation and emmetropization. Vision Research, 42(21), 2409–2417. https://doi.org/10.1016/S0042-6989(02)00262-6

Sivak, J. G., & Millodot, M. (1974). Axial chromatic aberration of eye with achromatizing lens. JOSA, 64(12), 1724–1725. https://doi.org/10.1364/JOSA.64.001724

Spitschan, M. (2019a). Melanopsin contributions to non-visual and visual function. Current Opinion in Behavioral Sciences, 30, 67–72. https://doi.org/10.1016/j.cobeha.2019.06.004

Spitschan, M. (2019b). Photoreceptor inputs to pupil control. Journal of Vision, 19(9), 5. https://doi.org/10.1167/19.9.5

Sravani, N. G., Nilagiri, V. K., & Bharadwaj, S. R. (2015). Photorefraction estimates of refractive power varies with the ethnic origin of human eyes. Scientific Reports, 5, 7976. https://doi.org/10.1038/srep07976

Stakenburg, M. (1991). Accommodation without pupillary constriction. Vision Research, 31(2), 267–273. https://doi.org/10.1016/0042-6989(91)90117-N

Stark, L. R., & Atchison, D. A. (1997). Pupil size, mean accommodation response and the fluctuations of accommodation. Ophthalmic & Physiological Optics: The Journal of the British College of Ophthalmic Opticians (Optometrists), 17(4), 316–323.

Stockman, A., Jägle, H., Pirzer, M., & Sharpe, L. T. (2008). The dependence of luminous efficiency on chromatic adaptation. Journal of Vision, 8(16), 1.1-26. https://doi.org/10.1167/8.16.1

Suchkov, N., Fernández, E. J., & Artal, P. (2019). Impact of longitudinal chromatic aberration on through-focus visual acuity. Optics Express, 27(24), 35935–35947. https://doi.org/10.1364/OE.27.035935

Thibos, L. N., Bradley, A., & López-Gil, N. (2013). Modelling the impact of spherical aberration on accommodation. Ophthalmic & Physiological Optics: The Journal of the British College of Ophthalmic Opticians (Optometrists), 33(4), 482–496. https://doi.org/10.1111/opo.12047

Thibos, L. N., Ye, M., Zhang, X., & Bradley, A. (1992). The chromatic eye: A new reduced-eye model of ocular chromatic aberration in humans. Applied Optics, 31(19), 3594–3600. Scopus. https://doi.org/10.1364/AO.31.003594

Tucker, J., & Charman, W. N. (1975). The depth-of-focus of the human eye for Snellen letters. American Journal of Optometry and Physiological Optics, 52(1), 3–21. https://doi.org/10.1097/00006324-197501000-00002

van der Wildt, G. J., & Bouman, M. A. (1971). An Accommodometer: An Apparatus for Measuring the Total Accommodation Response of the Human Eye. Applied Optics, 10(8), 1950–1958. https://doi.org/10.1364/AO.10.001950

Wald, G., & Griffin, D. R. (1947). The Change in Refractive Power of the Human Eye in Dim and Bright Light. JOSA, 37(5), 321–336. https://doi.org/10.1364/JOSA.37.000321

Wang, B., & Ciuffreda, K. J. (2006). Depth-of-focus of the human eye: Theory and clinical implications. Survey of Ophthalmology, 51(1), 75–85. https://doi.org/10.1016/j.survophthal.2005.11.003

Ward, P. A., & Charman, W. N. (1985). Effect of pupil size on steady state accommodation. Vision Research, 25(9), 1317–1326. https://doi.org/10.1016/0042-6989(85)90047-1

Wilson, D. (1973). A centre for accommodative vergence motor control. Vision Research, 13(12), 2491–2503. https://doi.org/10.1016/0042-6989(73)90246-0

Yao, P., Lin, H., Huang, J., Chu, R., & Jiang, B. (2010). Objective depth-of-focus is different from subjective depth-of-focus and correlated with accommodative microfluctuations. Vision Research, 50(13), 1266–1273. https://doi.org/10.1016/j.visres.2010.04.011

