## Supplementary Material for "Accommodation and wavelength: the effect of longitudinal chromatic aberration on the stimulus-response curve"

### Methods.

In this section we describe the methods used in three experiments where we measured the observers' accommodation response curve to different spectra. In experiments 1 and 2, the accommodation function was sampled by changing the physical distance of the stimuli, with the angular size of the diffuser changing concurrently in experiment 1 and being kept constant in experiment 2. In experiment 3, the accommodation function was measured by using trial lenses to simulate a larger range of optical distances, and the visual acuity of participants was measured concurrently. Experiments 1 and 2 used the same apparatus, thus, they are described together, while experiment 3 is described separately where necessary.

#### Participants.

In experiments 1 and 2, only participants that did not require visual correction (i.e., spectacles or contact lenses) to read or perform other daily activities were selected. The mean visual acuity of the sample was logMAR 0.03 with a range between logMAR -0.1 and 0.23. This means that the smallest characters they could read had a stroke width of 1.1 arcmin on average, with a range in the sample between 0.8 and 1.7 arcmin. In experiment 3, two of the ten participants normally used spectacles to read or perform activities at near distances (with corrections of approximately -0.7D and -2.5D) but performed the experiment without them, as they would change the intended accommodative demands.

#### Photorefractor calibration.

To find an individual correction factor for each participant we followed the method described by Sravani et al. (2015). A fixation stimulus illuminated by the green LED is presented at 1D from the participant and viewed monocularly through the opposite eye for which the calibration was being performed. The eye being calibrated (right eye in experiments 1 and 2, and left eye in experiment 3) was covered by an infrared filter, allowing to measure its refractive state while occluding the stimulus. A series of trial lenses from -4D to 7D in 1D steps were also placed in front of this eye, and refraction was measured binocularly for at least 30 seconds for each of the lenses. This method allows to obtain the defocus measured by the photorefractor for objective values of defocus introduced for the calibrated eye through the trial lenses, while also accounting for the changes in accommodation by concurrently measuring the refraction of the left eye that views the stimulus. An individual correction factor was obtained by plotting the average differences in refraction between the two eyes as a function of the trial lens used and fitting a linear regression through the linear portion of this function. The inverse of the resulting slope was then used to rescale all the refractive data obtained for this participant. An example of the results of the calibration procedure obtained for one subject are shown in Supp Fig 1. In experiment 3, to account for the differences in refractive error between the two eyes, the average refractive difference with no trial lens (0D) was obtained for each participant and subtracted from their data.

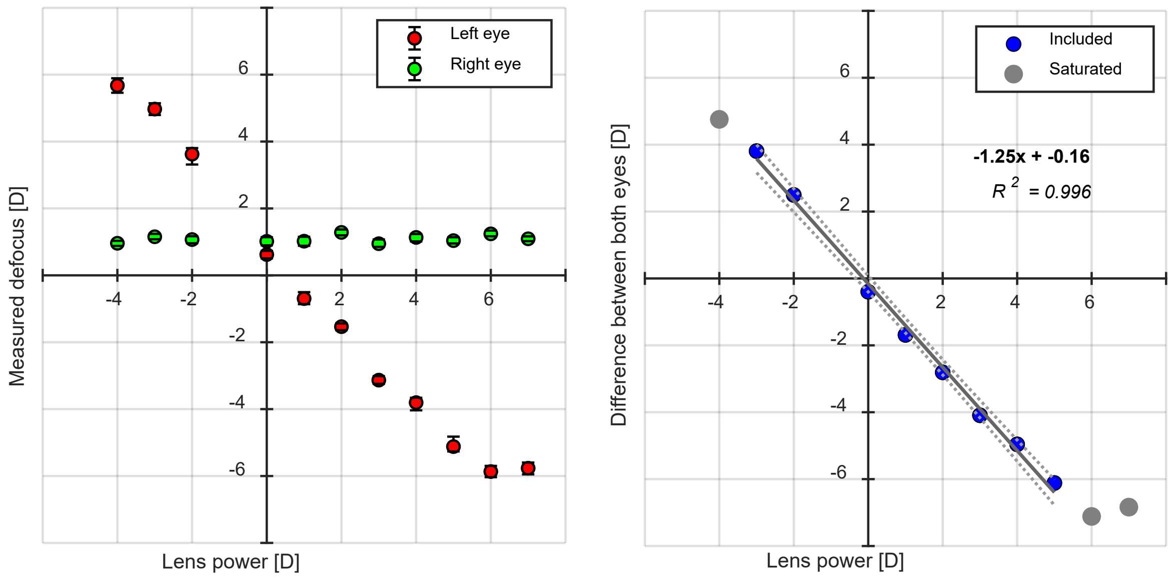


*Supp Fig 1. Results of the calibration procedure for one participant. The left panel shows the median and 25th and 75th percentiles of the measured defocus in both eyes as a function of the power of the lens used in front of the left eye. The right eye was uncovered and accommodating on a fixed target, while the left eye was covered by an infrared filter and different lenses. The right panel shows the difference in defocus between both eyes and the fitted linear regression. The steep slope indicates that the photorefractor overestimates the defocus in the left eye of this participant, measuring 1.25 dioptres for each 1 dioptre of real defocus. The inverse of this slope can be used to rescale the refraction measurements and correct the overestimation.*

### Results.

#### Determining the accommodative range of observers.

Accommodative range varies widely among individuals, particularly in a sample of observers with differences in age and refractive error. Within the accommodative range the response is expected to be quasi-linear with respect to the demand, while beyond it – that is, for demands higher than the near point of accommodation or lower than the far point – the response becomes saturated as the power of the crystalline lens cannot longer increase or decrease, respectively. The slope of the accommodation response curve is usually assessed within this linear accommodative range; thus, it was important to determine the near and far point of accommodation for each individual observer in our sample.

To do this, we calculated for each participant the gradient of the accommodation response as a function of distance in dioptres for each illuminant. At distances where the gradient dropped by 50% or more when compared to the overall median gradient, the response was determined to be saturated, while the distances where the gradient was maintained were taken to be within the accommodative range or linear portion of the accommodation response curve. This process was done for individual illuminants, such that the response to each could saturate at different distances (due to the differences in accommodative demand caused by LCA). The results of this process were visually inspected and agreed well with the evaluation of the experimenter (see Supp Fig 2 for an example of the results for one subject). In all the analyses presented in following sections, the saturated portion of the accommodation response was omitted (i.e., only accommodative and pupil responses within the linear portion of the accommodation response curve were included), except for section 2.3.6, where some analyses include responses beyond the accommodative range, as detailed there.

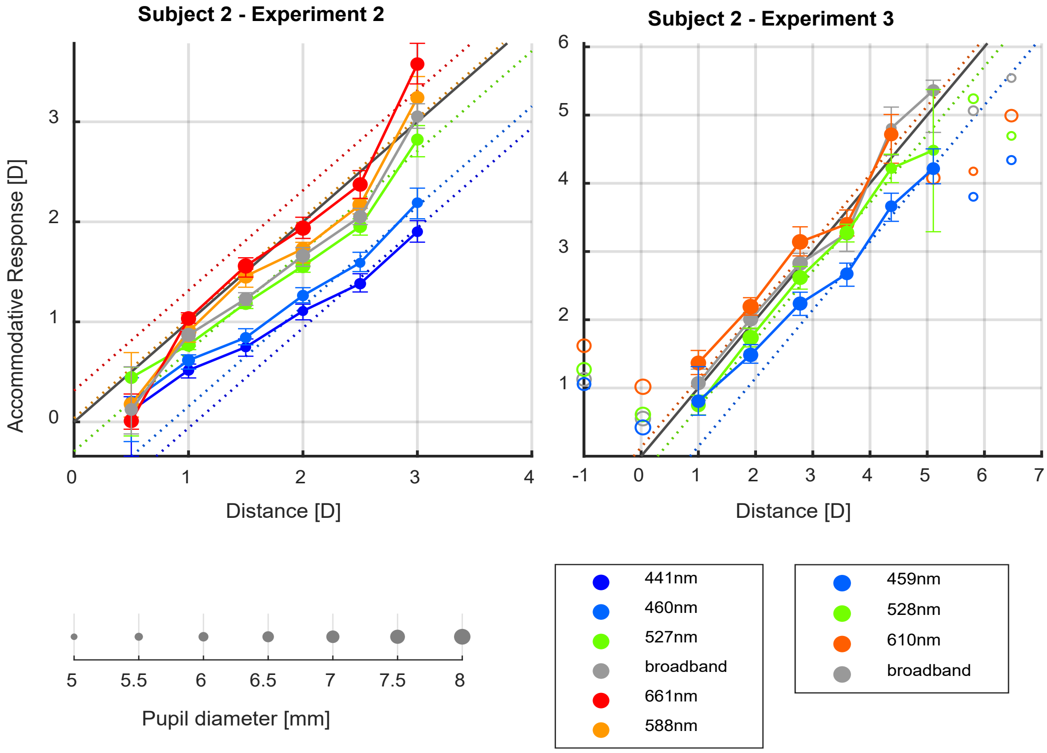


*Supp Fig 2. Median accommodation response as a function of distance in dioptres for one participant in experiments 2 and 3. The error bars represent the 25th and 75th percentiles of the response. The filled markers and continuous lines represent the portion of the response deemed to be within the accommodative range, while the unconnected open markers with no error bars are the saturated response. The colour of the markers represents the illuminant and the size the corresponding median pupil diameter, as indicated by the legend. The continuous black line represents the 1:1 ideal response, while the dotted coloured lines represent the 1:1 response corrected by the LCA defocus for each illuminant.*

Effects of LCA on the accommodation response curve.

| Experiment & Illuminant | | Slope [D/D] | | | | | | | Intercept [D] | | | |
|---|---|---|---|---|---|---|---|---|---|---|---|---|
| | | Est. | 95% CI | | RE SD | t-ratio | df | p-value | Est. | RE SD | t-ratio | p-value |
| 1 | 661nm | 0.89 | 0.81 | 0.97 | 0.10 | | | | 0.33 | 0.16 | | |
| | 588nm | 0.79 | 0.74 | 0.84 | 0.04 | -3.84 | 1246 | *<0.001* | 0.36 | 0.07 | 0.69 | 0.488 |
| | 527nm | 0.79 | 0.74 | 0.84 | 0.03 | -4.15 | 1246 | *<0.001* | 0.21 | 0.08 | -2.05 | *0.041* |
| | 460nm | 0.71 | 0.65 | 0.77 | 0.06 | -5.90 | 1246 | *<0.001* | 0.19 | 0.19 | -1.61 | 0.107 |
| | 441nm | 0.63 | 0.58 | 0.69 | 0.05 | -9.07 | 1246 | *<0.001* | 0.23 | 0.24 | -0.95 | 0.341 |
| | broadband | 0.90 | 0.84 | 0.95 | 0.05 | 0.08 | 1246 | 0.940 | 0.12 | 0.09 | -3.67 | *<0.001* |
| 2 | 661nm | 1.07 | 0.91 | 1.22 | 0.23 | | | | -0.16 | 0.71 | | |
| | 588nm | 0.94 | 0.88 | 1.01 | 0.07 | -3.89 | 3157 | *<0.001* | -0.08 | 0.10 | 1.43 | 0.154 |
| | 527nm | 0.92 | 0.84 | 1.00 | 0.10 | -3.66 | 3157 | *<0.001* | -0.21 | 0.23 | -0.53 | 0.595 |
| | 460nm | 0.82 | 0.72 | 0.91 | 0.13 | -5.07 | 3157 | *<0.001* | -0.21 | 0.30 | -0.46 | 0.648 |
| | 441nm | 0.78 | 0.69 | 0.86 | 0.11 | -6.59 | 3157 | *<0.001* | -0.21 | 0.26 | -0.49 | 0.626 |

| | | | | | | | | | | | | |
|---|---|---|---|---|---|---|---|---|---|---|---|---|
| | broadband | 1.00 | 0.93 | 1.07 | 0.08 | -1.86 | 3157 | 0.063 | -0.30 | 0.15 | -2.02 | *0.044* |
| **3** | 610nm | 1.15 | 0.67 | 1.64 | 0.65 | | | | -1.16 | 3.24 | -0.95 | |
| | 528nm | 1.09 | 0.81 | 1.36 | 0.36 | -0.49 | 1774 | 0.624 | -1.13 | 1.64 | 0.06 | 0.956 |
| | 459nm | 0.95 | 0.66 | 1.24 | 0.38 | -1.40 | 1774 | 0.161 | -0.93 | 1.84 | 0.33 | 0.744 |
| | broadband | 1.11 | 0.92 | 1.31 | 0.26 | -0.43 | 1774 | 0.667 | -0.98 | 1.24 | 0.38 | 0.702 |

*Supp Table 1. Linear mixed models of accommodation as a function of distance in dioptres and illuminant for each experiment: Accommodation ~ Distance * Illuminant + (1 + Distance * Illuminant | ID). The estimated coefficients and their 95% confidence intervals (95% CI) have been used to calculate the estimated slopes (in dioptres/dioptres) and intercepts (in dioptres) for each illuminant. The intercept specifies where the average person accommodates for a stimulus at infinity under the specified illuminant, and the slope specifies how that changes with dioptric distance. The random effects standard deviations (RE SD), t-ratios, degrees of freedom (df), and p-values are also shown. The t-tests compare within each experiment, the slope and intercept estimates of each illuminant with the estimates for the longest-wavelength illuminant.*

Variability in the accommodation responses to narrowband and broadband illuminants.

| **Experiment** | **Parameter** | **Estimate** | **CI 95%** | | **t-test** | **df** | **p-value** | **RE SD** |
|---|---|---|---|---|---|---|---|---|
| 1 & 2 | Intercept | 0.155 | 0.12 | 0.19 | 8.94 | 4420 | *<0.001* | 0.06 |
| | 460nm | -0.008 | -0.02 | 0.00 | -2.06 | 4420 | *0.039* | 0.01 |
| | 527nm | -0.023 | -0.03 | -0.01 | -5.35 | 4420 | *<0.001* | 0.01 |
| | 588nm | -0.050 | -0.07 | -0.03 | -6.18 | 4420 | *<0.001* | 0.03 |
| | 661nm | -0.043 | -0.05 | -0.03 | -7.22 | 4420 | *<0.001* | 0.02 |
| | broadband | -0.022 | -0.03 | -0.01 | -4.63 | 4420 | *<0.001* | 0.01 |
| | Refraction | 0.020 | 0.01 | 0.03 | 4.21 | 4420 | *<0.001* | 0.02 |
| 3 | Intercept | 0.134 | 0.10 | 0.16 | 8.67 | 2190 | *<0.001* | 0.05 |
| | 528nm | -0.028 | -0.04 | -0.01 | -3.78 | 2190 | *<0.001* | 0.02 |
| | 610nm | -0.031 | -0.05 | -0.02 | -3.93 | 2190 | *<0.001* | 0.02 |
| | broadband | -0.020 | -0.03 | -0.01 | -2.81 | 2190 | *0.005* | 0.02 |
| | Refraction | 0.031 | 0.02 | 0.04 | 6.45 | 2190 | *<0.001* | 0.01 |

*Supp Table 2. Linear mixed model results of the root-mean-square errors (RMSEs) of an unconstrained linear fit through the within-trial accommodation response, as a function of refraction and illuminant, in experiments 1 and 2, and experiment 3. Coefficient estimates and their 95% confidence interval (CI 95%) are shown, as well as degrees of freedom (df), t-ratios p-values, and the random effects standard deviation (RE SD). The intercepts of the models are the corresponding shortest wavelength illuminants (441nm in experiments 1 and 2, and 459nm in experiment 3) at zero dioptres of refraction.*

Accommodation and pupil size.

| **Experiment** | **Parameters** | **Estimate** | **CI 95%** | | **t-ratio** | **df** | **p-value** |
|---|---|---|---|---|---|---|---|
| 1 | Intercept | 6.11 | 5.79 | 6.42 | 38.09 | 1448 | <0.001 |
| | 460nm | 0.22 | 0.11 | 0.33 | 3.95 | 1448 | <0.001 |

| | | | | | | | |
|---|---|---|---|---|---|---|---|
| | 527nm | 0.97 | 0.67 | 1.27 | 6.34 | 1448 | <0.001 |
| | 588nm | 1.66 | 1.39 | 1.94 | 11.98 | 1448 | <0.001 |
| | 661nm | 1.70 | 1.38 | 2.01 | 10.57 | 1448 | <0.001 |
| | broadband | 0.86 | 0.62 | 1.10 | 7.12 | 1448 | <0.001 |
| | Refraction | -0.75 | -0.89 | -0.62 | -10.81 | 1448 | <0.001 |
| 2 | Intercept | 5.28 | 4.80 | 5.76 | 21.68 | 3494 | <0.001 |
| | 460nm | 0.23 | 0.16 | 0.30 | 6.22 | 3494 | <0.001 |
| | 527nm | 0.75 | 0.59 | 0.90 | 9.48 | 3494 | <0.001 |
| | 588nm | 1.40 | 1.21 | 1.58 | 14.67 | 3494 | <0.001 |
| | 661nm | 1.26 | 1.11 | 1.41 | 16.22 | 3494 | <0.001 |
| | broadband | 0.70 | 0.55 | 0.86 | 8.82 | 3494 | <0.001 |
| | Refraction | -0.16 | -0.26 | -0.06 | -3.12 | 3494 | 0.002 |
| 3 | Intercept | 5.63 | 5.12 | 6.15 | 21.62 | 912 | <0.001 |
| | 528nm | 0.30 | 0.18 | 0.42 | 4.81 | 912 | <0.001 |
| | 610nm | 0.45 | 0.30 | 0.59 | 6.06 | 912 | <0.001 |
| | broadband | 0.23 | 0.14 | 0.31 | 5.18 | 912 | <0.001 |
| | Refraction | -0.18 | -0.29 | -0.06 | -3.02 | 912 | 0.003 |

*Supp Table 3. Linear mixed model results of pupil diameter for experiments 1, 2 and 3, fitting Pupil Diameter ~ Accommodation + Illuminant + (1 + Accommodation + Illuminant | ID). Coefficient estimates, their 95% confidence interval (CI 95%) and standard errors (SE) are shown, as well as degrees of freedom (df), t-ratios and p-values. The intercepts of the models are the corresponding shortest wavelength illuminants (441nm in experiments 1 and 2, and 459nm in experiment 3) at zero dioptres of refraction.*

Accommodative error and visual acuity.

| **Visual acuity [logMAR] - Linear portion of the accommodation function** | | | | | | | |
|---|---|---|---|---|---|---|---|
| **Parameters** | **Estimate** | **CI 95%** | | **t-ratio** | **df** | **p-value** | **RE SD** |
| Intercept | 0.04 | -0.06 | 0.15 | 0.79 | 8.29 | 0.452 | 0.16 |
| Error magnitude [D] | 0.10 | -0.02 | 0.21 | 1.69 | 8.66 | 0.126 | 0.17 |
| Positive sign | 0.01 | -0.07 | 0.09 | 0.26 | 7.04 | 0.800 | 0.11 |
| Illuminant: 528nm | ***-0.11*** | ***-0.17*** | ***-0.06*** | ***-3.95*** | ***8.72*** | ***0.004*** | ***0.08*** |
| Illuminant: 610nm | ***-0.08*** | ***-0.14*** | ***-0.02*** | ***-2.72*** | ***8.70*** | ***0.024*** | ***0.08*** |
| Illuminant: broadband | ***-0.09*** | ***-0.12*** | ***-0.05*** | ***-4.53*** | ***9.09*** | ***0.001*** | ***0.05*** |
| Error magnitude [D] * Positive sign | 0.02 | -0.10 | 0.14 | 0.31 | 9.70 | 0.761 | 0.16 |

*Supp Table 4. Linear mixed model results of visual acuity over the linear portion of the accommodation response curve, as a function of illuminant, accommodative error magnitude, accommodative error sign, and their interaction. The coefficient estimates, their 95%*

*confidence intervals (CI 95%), t-ratios, degrees of freedom (df), p-values, and random effects standard deviations (RE SD) are shown.*

| **Visual acuity [logMAR] - Overaccommodation** | | | | | | | |
|---|---|---|---|---|---|---|---|
| **Parameters** | **Estimate** | **CI 95%** | | **df** | **t-ratio** | **p-value** | **RE SD** |
| Intercept<br>*(4mm, 0D, 459nm)* | 0.13 | -0.06 | 0.32 | 4.52 | 1.31 | 0.252 | 0.29 |
| Error magnitude [D] | 0.21 | 0.10 | 0.31 | 4.38 | 3.84 | 0.016 | 0.14 |
| Pupil Diameter [mm] | -0.07 | -0.15 | 0.01 | 5.80 | -1.80 | 0.123 | 0.10 |
| Illuminant: 528nm | -0.05 | -0.11 | 0.01 | 10.09 | -1.67 | 0.126 | 0.08 |
| Illuminant: 610nm | -0.01 | -0.08 | 0.06 | 9.62 | -0.23 | 0.822 | 0.09 |
| Illuminant: broadband | -0.04 | -0.09 | 0.01 | 13.18 | -1.68 | 0.116 | 0.05 |
| Error magnitude [D] * Pupil Diameter [mm] | ***0.08*** | ***0.02*** | ***0.13*** | ***5.39*** | ***2.78*** | ***0.036*** | ***0.07*** |
| **Visual acuity [logMAR] - Underaccommodation** | | | | | | | |
| **Parameters** | **Estimate** | **CI 95%** | | **df** | **t-ratio** | **p-value** | **RE SD** |
| Intercept<br>*(4mm, 0D, 459nm)* | 0.14 | 0.04 | 0.25 | 5.08 | 2.61 | *0.047* | 0.14 |
| Error magnitude [D] | 0.06 | 0.00 | 0.12 | 9.97 | 1.82 | 0.098 | 0.07 |
| Pupil Diameter [mm] | -0.05 | -0.12 | 0.02 | 7.09 | -1.44 | 0.192 | 0.09 |
| Illuminant: 528nm | -0.07 | -0.14 | -0.01 | 7.31 | -2.16 | 0.066 | 0.08 |
| Illuminant: 610nm | -0.04 | -0.11 | 0.03 | 7.48 | -1.21 | 0.264 | 0.09 |
| Illuminant: broadband | -0.07 | -0.14 | -0.01 | 4.83 | -2.15 | 0.086 | 0.08 |
| Error magnitude [D] * Pupil Diameter [mm] | 0.04 | -0.02 | 0.11 | 3.53 | 1.31 | 0.268 | 0.09 |

*Supp Table 5. Linear mixed models' results of visual acuity for positive accommodative errors (top) and negative accommodative errors (bottom), as a function of error magnitude, pupil diameter, their interaction, and illuminant. The coefficient estimates, their 95% confidence intervals (CI 95%), t-ratios, degrees of freedom (df), p-values, and random effects standard deviations (RE SD) are shown.*

### Accommodation response curves for individual participants.

In this section, we present the accommodation response curves of individual participants in experiments 1, 2 and 3. The figures present the accommodation response as a function of the distance in dioptres.

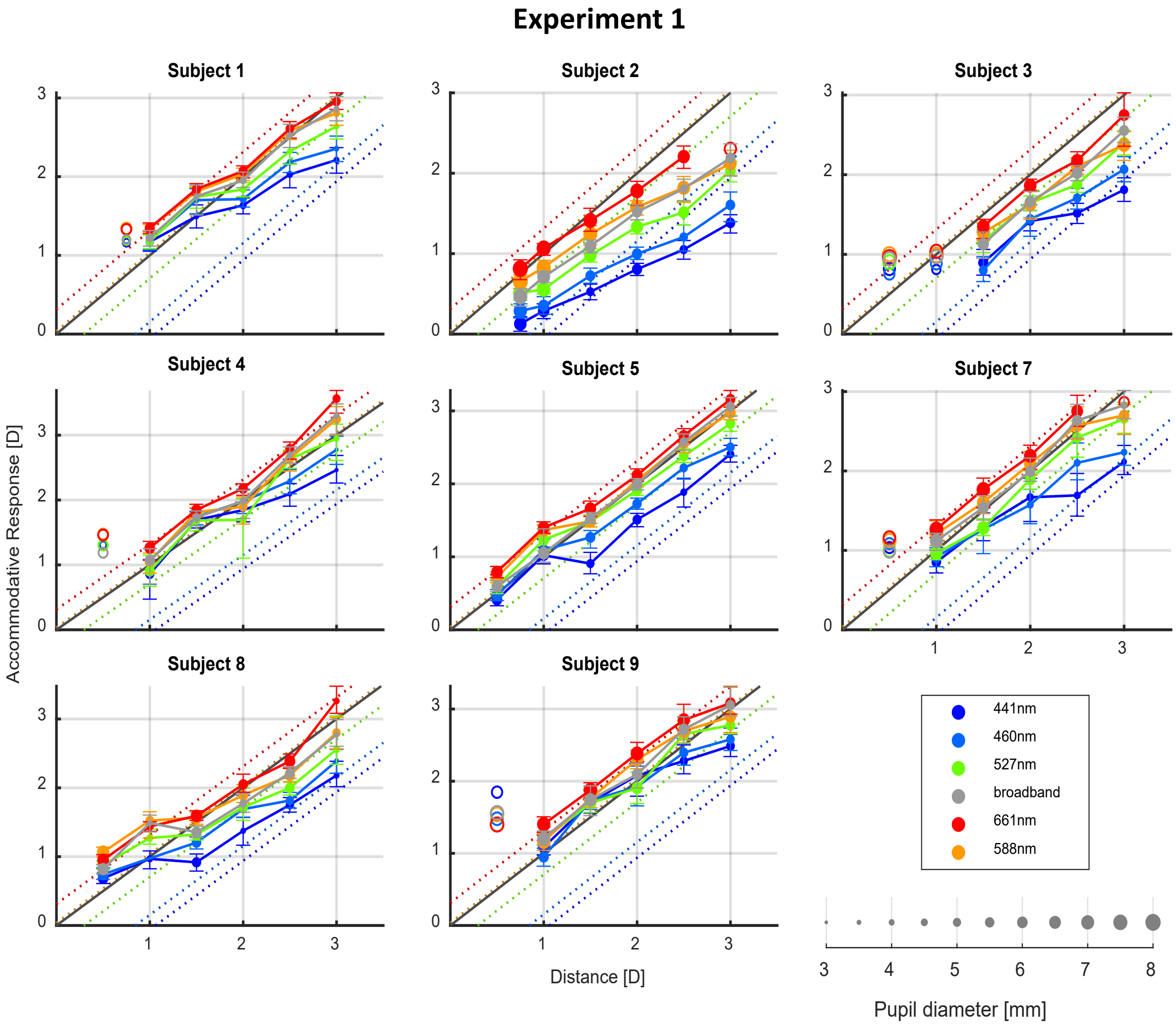


*Supp Fig 3. Median accommodation response as a function of distance for individual participants of experiment 1. Each colour represents one illuminant, and the marker sizes represent the median pupil size for each distance and illuminant. The error bars represent the 25$^{th}$ and 75$^{th}$ percentiles of the accommodation response. The filled markers and continuous lines represent the portion of the response curve deemed to be linear, while the unconnected open markers with no error bars represent the portions of the curve identified as saturated. The continuous grey line represents the one-to-one response, while the dotted coloured lines represent the one-to-one response corrected by the LCA defocus for the peak wavelength of each illuminant.*

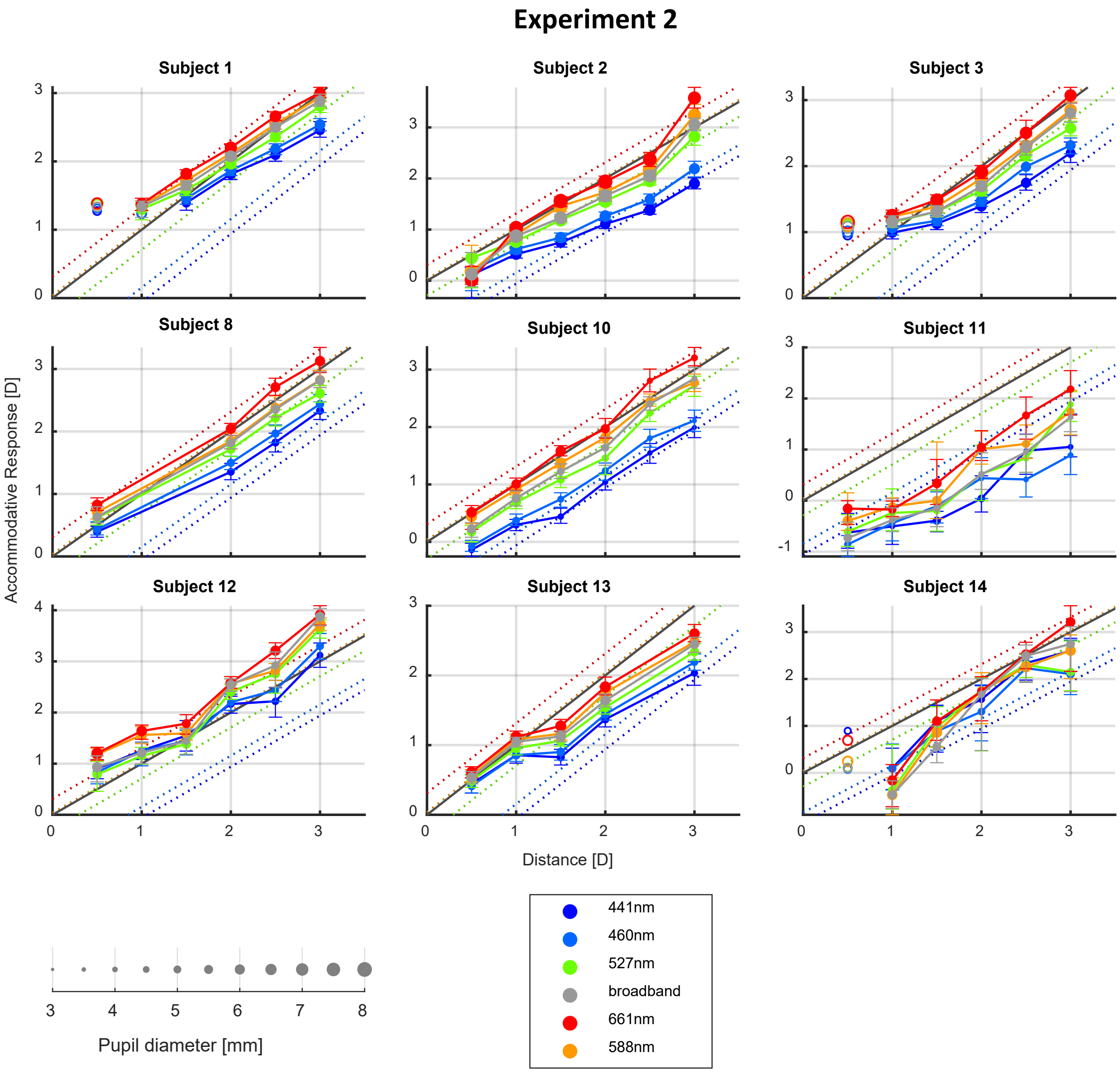


*Supp Fig 4. Median accommodation response as a function of distance for individual participants of experiment 2. Each colour represents one illuminant, and the marker sizes represent the median pupil size for each distance and illuminant. The error bars represent the 25th and 75th percentiles of the accommodation response. The filled markers and continuous lines represent the portion of the response curve deemed to be linear, while the unconnected open markers with no error bars represent the portions of the curve identified as saturated. The continuous grey line represents the one-to-one response, while the dotted coloured lines represent the one-to-one response corrected by the LCA defocus for the peak wavelength of each illuminant.*

### Experiment 3

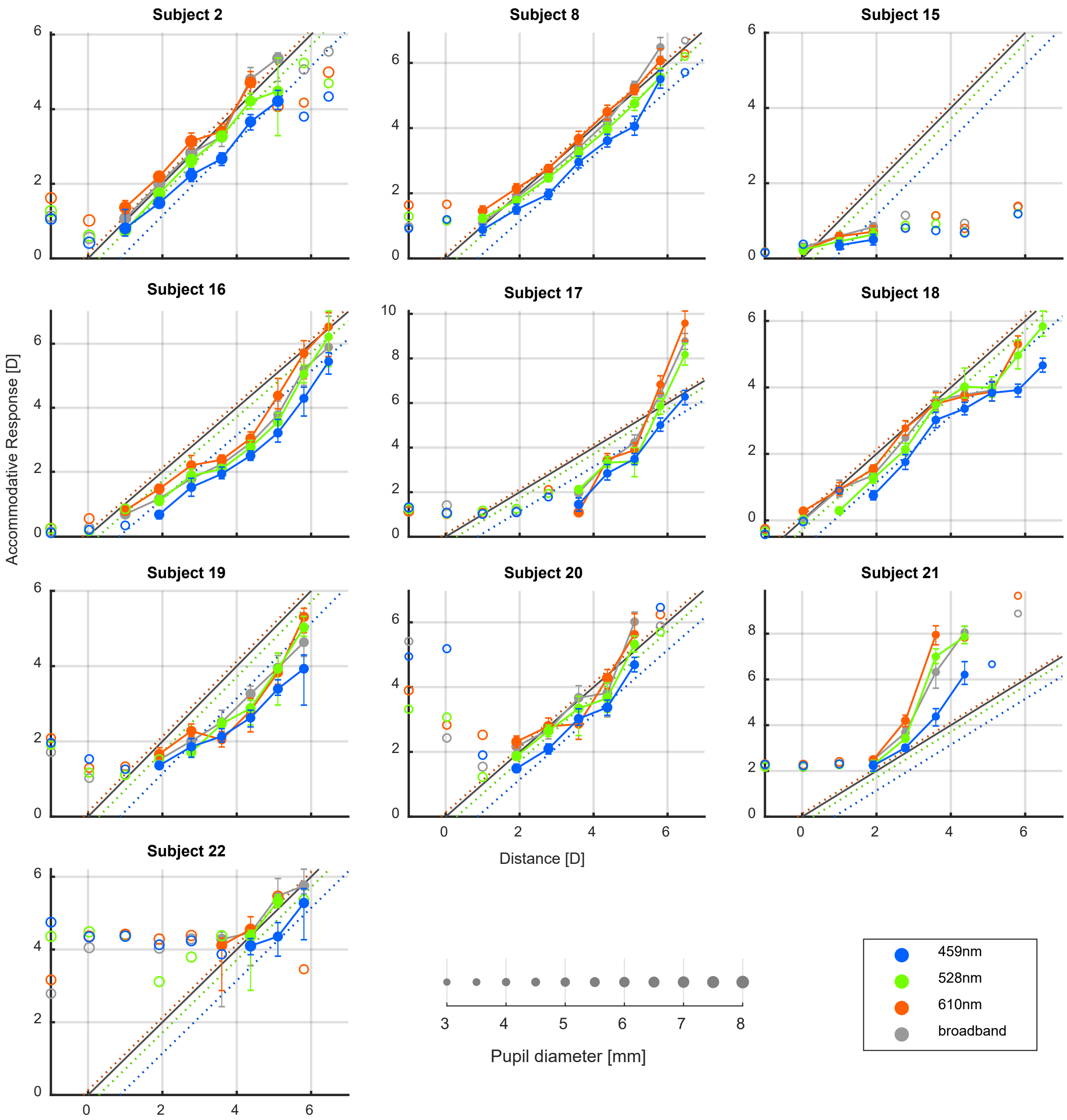


*Supp Fig 5. Median accommodation response as a function of distance for individual participants of experiment 3. Each colour represents one illuminant, and the marker sizes represent the median pupil size for each distance and illuminant. The error bars represent the 25th and 75th percentiles of the accommodation response. The filled markers and continuous lines represent the portion of the response curve deemed to be linear, while the unconnected open markers with no error bars represent the portions of the curve identified as saturated. The continuous grey line represents the one-to-one response, while the dotted coloured lines represent the one-to-one response corrected by the LCA defocus for the peak wavelength of each illuminant.*

Results of the linear mixed models of accommodation for individual participants.

In this section, we present additional results of the linear mixed models used on the linear portion of the accommodation response curves, as described in the manuscript. The table shows the slopes and intercepts estimated for individual participants under each illuminant, which were calculated from the estimated random effects coefficients of the linear mixed models fitted to the data of each experiment.

| subject | Experiment 1 | | | | | |
|---|---|---|---|---|---|---|
| | broadband | 661 nm | 588 nm | 527 nm | 460 nm | 441 nm |
| 1 | 0.89 (0.02)<br>*0.19 (0.02)* | 0.81 (0.02)<br>*0.55 (0.03)* | 0.81 (0.02)<br>*0.43 (0.03)* | 0.79 (0.01)<br>*0.30 (0.03)* | 0.68 (0.02)<br>*0.37 (0.05)* | 0.62 (0.02)<br>*0.41 (0.06)* |
| 2 | 0.85 (0.02)<br>*0.06 (0.02)* | 0.78 (0.02)<br>*0.22 (0.03)* | 0.73 (0.02)<br>*0.35 (0.03)* | 0.78 (0.01)<br>*0.06 (0.03)* | 0.70 (0.03)<br>*-0.09 (0.04)* | 0.65 (0.02)<br>*-0.17 (0.05)* |
| 3 | 0.97 (0.02)<br>*-0.05 (0.03)* | 0.85 (0.03)<br>*0.12 (0.05)* | 0.81 (0.02)<br>*0.33 (0.04)* | 0.83 (0.02)<br>*0.18 (0.03)* | 0.82 (0.03)<br>*-0.05 (0.06)* | 0.64 (0.03)<br>*0.27 (0.08)* |
| 4 | 0.90 (0.02)<br>*0.12 (0.03)* | 1.10 (0.02)<br>*0.14 (0.04)* | 0.82 (0.02)<br>*0.26 (0.03)* | 0.77 (0.02)<br>*0.23 (0.03)* | 0.67 (0.03)<br>*0.38 (0.06)* | 0.59 (0.03)<br>*0.46 (0.08)* |
| 5 | 0.92 (0.02)<br>*0.12 (0.02)* | 0.94 (0.02)<br>*0.33 (0.03)* | 0.82 (0.02)<br>*0.32 (0.03)* | 0.80 (0.02)<br>*0.24 (0.03)* | 0.75 (0.03)<br>*0.14 (0.05)* | 0.70 (0.02)<br>*0.05 (0.06)* |
| 7 | 0.87 (0.02)<br>*0.16 (0.03)* | 0.93 (0.03)<br>*0.34 (0.04)* | 0.76 (0.02)<br>*0.40 (0.03)* | 0.80 (0.02)<br>*0.16 (0.03)* | 0.68 (0.03)<br>*0.18 (0.05)* | 0.60 (0.03)<br>*0.25 (0.07)* |
| 8 | 0.85 (0.02)<br>*0.21 (0.03)* | 0.80 (0.02)<br>*0.52 (0.03)* | 0.77 (0.02)<br>*0.44 (0.03)* | 0.77 (0.02)<br>*0.26 (0.03)* | 0.68 (0.03)<br>*0.24 (0.05)* | 0.65 (0.03)<br>*0.12 (0.07)* |
| 9 | 0.91 (0.02)<br>*0.17 (0.03)* | 0.95 (0.02)<br>*0.39 (0.04)* | 0.82 (0.02)<br>*0.37 (0.03)* | 0.79 (0.02)<br>*0.28 (0.03)* | 0.70 (0.03)<br>*0.37 (0.05)* | 0.60 (0.03)<br>*0.49 (0.07)* |
| | **Experiment 2** | | | | | |
| | broadband | 661 nm | 588 nm | 527 nm | 460 nm | 441 nm |
| 1 | 1.10 (0.02)<br>*-0.28 (0.05)* | 0.84 (0.02)<br>*0.55 (0.05)* | 1.00 (0.01)<br>*-0.19 (0.02)* | 0.99 (0.03)<br>*-0.28 (0.07)* | 0.95 (0.03)<br>*-0.33 (0.08)* | 0.89 (0.03)<br>*-0.30 (0.08)* |
| 2 | 0.89 (0.02)<br>*-0.10 (0.06)* | 1.20 (0.03)<br>*-0.41 (0.07)* | 0.92 (0.01)<br>*0.00 (0.03)* | 0.79 (0.04)<br>*-0.04 (0.09)* | 0.62 (0.04)<br>*-0.04 (0.09)* | 0.56 (0.04)<br>*-0.11 (0.09)* |
| 3 | 0.99 (0.02)<br>*-0.18 (0.06)* | 0.90 (0.03)<br>*0.24 (0.07)* | 1.00 (0.01)<br>*-0.13 (0.03)* | 0.92 (0.04)<br>*-0.14 (0.09)* | 0.86 (0.04)<br>*-0.20 (0.09)* | 0.81 (0.04)<br>*-0.19 (0.09)* |
| 8 | 1.00 (0.02)<br>*-0.27 (0.06)* | 0.92 (0.03)<br>*0.30 (0.06)* | 0.98 (0.01)<br>*-0.17 (0.03)* | 0.96 (0.04)<br>*-0.30 (0.08)* | 0.91 (0.04)<br>*-0.42 (0.08)* | 0.87 (0.04)<br>*-0.46 (0.08)* |
| 10 | 0.96 (0.02)<br>*-0.31 (0.05)* | 1.20 (0.03)<br>*-0.23 (0.05)* | 0.89 (0.01)<br>*-0.05 (0.03)* | 0.92 (0.03)<br>*-0.37 (0.07)* | 0.78 (0.03)<br>*-0.45 (0.07)* | 0.72 (0.04)<br>*-0.50 (0.08)* |

| | | | | | | |
|---|---|---|---|---|---|---|
| 11 | 0.92 (0.03) | 1.10 (0.03) | 0.82 (0.01) | 0.90 (0.04) | 0.76 (0.04) | 0.74 (0.04) |
| | *-0.30 (0.06)* | *-1.10 (0.06)* | *0.12 (0.03)* | *-0.26 (0.09)* | *-0.30 (0.09)* | *-0.26 (0.09)* |
| 12 | 1.10 (0.02) | 1.10 (0.03) | 0.99 (0.01) | 1.10 (0.04) | 0.91 (0.04) | 0.77 (0.04) |
| | *-0.56 (0.06)* | *0.39 (0.07)* | *-0.13 (0.03)* | *-0.51 (0.09)* | *-0.35 (0.09)* | *-0.17 (0.09)* |
| 13 | 1.00 (0.02) | 0.81 (0.03) | 0.99 (0.01) | 0.96 (0.03) | 0.93 (0.03) | 0.90 (0.04) |
| | *-0.23 (0.05)* | *0.21 (0.05)* | *-0.13 (0.03)* | *-0.22 (0.07)* | *-0.30 (0.07)* | *-0.28 (0.07)* |
| 14 | 1.00 (0.03) | 1.50 (0.03) | 0.90 (0.01) | 0.76 (0.04) | 0.63 (0.04) | 0.72 (0.04) |
| | *-0.44 (0.06)* | *-1.50 (0.07)* | *-0.09 (0.03)* | *0.22 (0.10)* | *0.47 (0.09)* | *0.35 (0.09)* |
| | **Experiment 3** | | | | | |
| | broadband | 610 nm | | 528 nm | 459 nm | |
| 2 | 1.00 (0.01) | 0.91 (0.03) | | 0.92 (0.03) | 0.82 (0.03) | |
| | *-1.60 (0.08)* | *0.44 (0.11)* | | *-1.60 (0.10)* | *-1.60 (0.15)* | |
| 8 | 1.10 (0.01) | 0.96 (0.02) | | 0.92 (0.02) | 0.89 (0.03) | |
| | *-1.60 (0.07)* | *0.29 (0.10)* | | *-1.40 (0.08)* | *-1.70 (0.14)* | |
| 16 | 0.98 (0.01) | 0.99 (0.03) | | 1.10 (0.02) | 0.95 (0.04) | |
| | *-1.40 (0.08)* | *-0.66 (0.11)* | | *-1.80 (0.09)* | *-1.80 (0.20)* | |
| 17 | 2.00 (0.03) | 2.60 (0.06) | | 1.80 (0.03) | 1.60 (0.07) | |
| | *1.80 (0.16)* | *-8.50 (0.31)* | | *2.50 (0.17)* | *3.20 (0.33)* | |
| 18 | 0.87 (0.01) | 0.82 (0.03) | | 0.96 (0.02) | 0.85 (0.05) | |
| | *-1.40 (0.09)* | *0.22 (0.11)* | | *-1.80 (0.11)* | *-1.90 (0.22)* | |
| 19 | 0.83 (0.01) | 0.82 (0.03) | | 1.10 (0.02) | 0.64 (0.04) | |
| | *-1.20 (0.09)* | *-0.18 (0.14)* | | *-2.40 (0.09)* | *-1.00 (0.18)* | |
| 20 | 1.00 (0.01) | 0.95 (0.04) | | 0.92 (0.03) | 0.91 (0.04) | |
| | *-1.50 (0.08)* | *0.20 (0.14)* | | *-1.30 (0.11)* | *-1.70 (0.18)* | |

*Supp Table 6. Individual slopes and intercepts estimated for the linear portion of the accommodation response curves of each participant to each illuminant used in experiments 1, 2 and 3. For each participant, the first row represents the estimated slopes for each illuminant, and the corresponding conditional standard deviations are shown between parentheses; and the second row in italics represents the estimated intercepts for each illuminant and the corresponding conditional standard deviations between parentheses.*

Visual acuity results for individual participants.

In this section, visual acuity as a function of accommodative error is plotted for individual participants, as described in the manuscript.

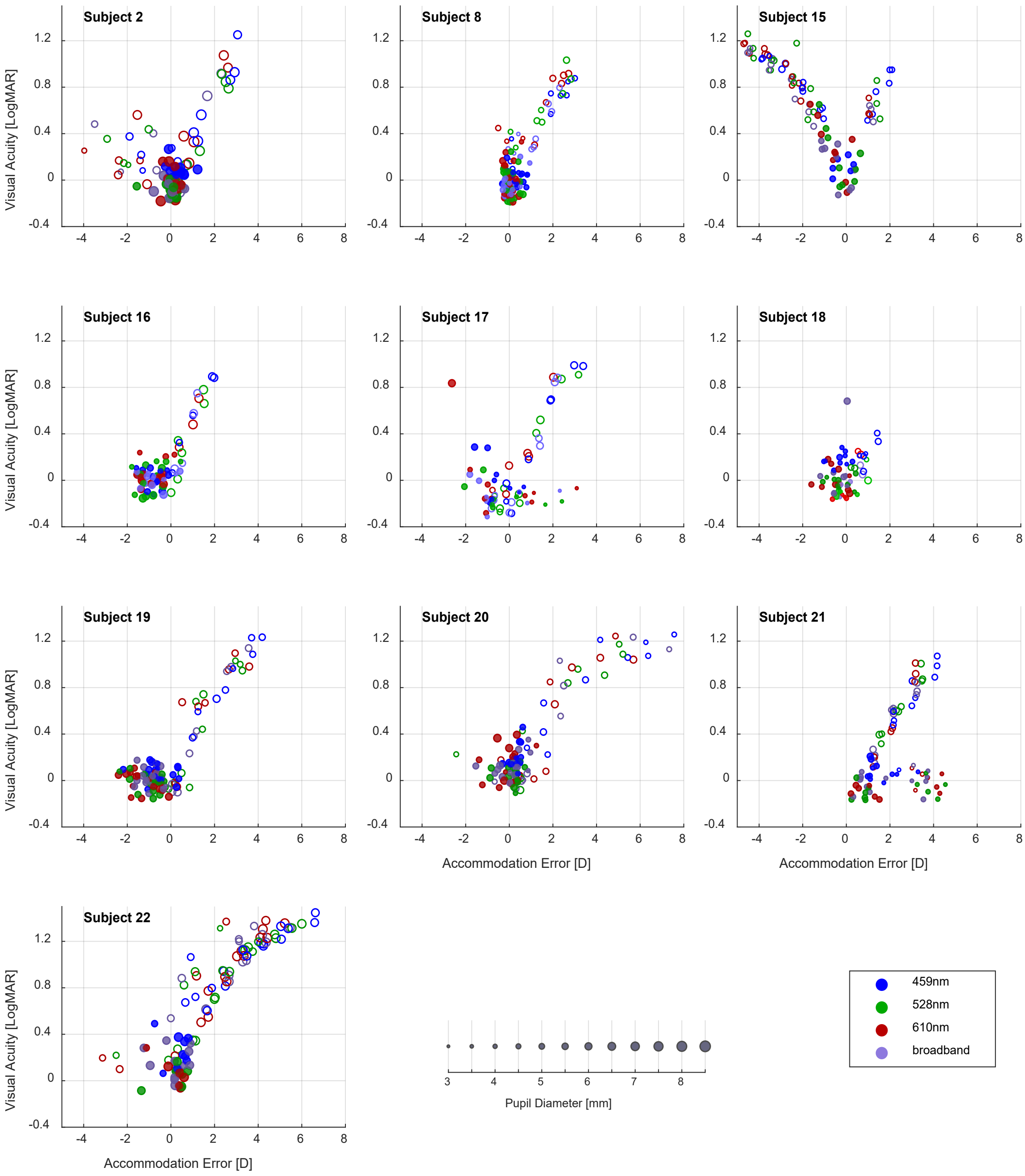


*Supp Fig 6. Visual acuity (VA) as a function of the median accommodative error for individual participants of experiment 3. Accommodative error is calculated as the accommodative demand subtracted from the accommodative response. Filled markers correspond to VA measurements over the linear portion of the accommodation response curve, while open markers correspond to measurements at distances where the accommodation function was saturated. The marker colours represent the illuminant used, and the marker sizes the corresponding median pupil size.*
